# A versatile tiling light sheet microscope for cleared tissues imaging

**DOI:** 10.1101/829267

**Authors:** Yanlu Chen, Xiaoliang Li, Dongdong Zhang, Chunhui Wang, Ruili Feng, Xuzhao Li, Yao Wen, Hao Xu, Xinyi Shirley Zhang, Xiao Yang, Yongyi Chen, Yi Feng, Bo Zhou, Bi-Chang Chen, Kai Lei, Shang Cai, Jie-Min Jia, Liang Gao

**Affiliations:** Key Laboratory of Structural Biology of Zhejiang Province, School of Life Sciences, Westlake University, Hangzhou, Zhejiang Province, 310024, China; Institute of Basic Medical Sciences, Westlake Institute for Advanced Study, Hangzhou, Zhejiang Province, 310024, China; Key Laboratory of Growth Regulation and Translation Research of Zhejiang Province, School of Life Sciences, Westlake University, Hangzhou, Zhejiang Province, 310024, China; Institute of Biology, Westlake Institute for Advanced Study, Hangzhou, Zhejiang Province, 310024, China; Nuohai Life Science Co.,Ltd, Shanghai, 201619, China; State Key Laboratory of Cell Biology, Shanghai Institute of Biochemistry and Cell Biology, Chinese Academy of Sciences, University of Chinese Academy of Sciences, Shanghai, 200031, China; State Key Laboratory of Proteomics, Beijing Proteome Research Center, National Center for Protein Sciences, Beijing Institute of Lifeomics, Beijing, 102206, China; Department of Clinical laboratory, Zhejiang Cancer Hospital, Hangzhou, Zhejiang Province, 310000, China; Department of Integrative Medicine and Neurobiology, School of Basic Medical Sciences, Fudan University, Shanghai, 200032, China; Research Center for Applied Sciences, Academia Sinica, Taipei, 11529, Taiwan

## Abstract

We present a tiling light sheet microscope compatible with all tissue clearing methods for rapid multicolor 3D imaging of centimeter-scale cleared tissues with micron-scale (4×4×10 μm^3^) to nanometer-scale (70×70×200 nm^3^) spatial resolution. The microscope uses tiling light sheets to achieve a more advanced multicolor 3D imaging ability than conventional light sheet microscopes. The illumination light is phase modulated to adjust the 3D imaging ability of the microscope based on the tissue size and the desired spatial resolution and imaging speed, and also to keep the microscope aligned. The ability of the microscope to align via phase modulation alone also makes the microscope reliable and easy to operate. Here we describe the working principle and design of the microscope. We demonstrate its imaging ability by imaging various cleared tissues prepared by different tissue clearing and tissue expansion techniques.

## Introduction

High-resolution 3D fluorescence imaging of biological tissues bridges the study of gene expression, cell morphology and cell distribution in tissues at subcellular, cellular, and tissue levels. It has mainly been performed by imaging physically sectioned tissue slices due to the opaqueness of biological tissues (Avants et al., 2011; Economo et al., 2016; Li et al., 2010; Micheva and Smith, 2007; Moffitt et al., 2018; Osten and Margrie, 2013, Ragan et al., 2012; Wang et al.,2020). Although tissue structures can be visualized in 3D with micron-scale or possibly higher spatial resolution by cutting and imaging a series of thin tissue slices, the method suffers from extremely low efficiency and many technical challenges in tissue sectioning, labeling, imaging and image registration processes. Not to mention structure distortions caused by the destructive tissue sectioning process, which hinders accurate registration and reconstruction of the imaged tissue slices to acquire the 3D image of the entire tissue.

Tissue clearing techniques enable the use of the latest 3D fluorescence imaging techniques to visualize tissue structures by making biological tissues transparent (Chen et al., 2017; Chung et al., 2013; Costantini et al., 2015; Economo et al., 2016; Epp et al., 2015; Erturk et al., 2012; Hama et al., 2011; Jing et al., 2018; Kurihara et al., 2015; Li et al., 2017; Pan et al., 2016; Seiriki et al., 2017; Susaki et al., 2014; Ueda et al., 2020). Among all 3D fluorescence imaging techniques, light sheet microscopy (LSM) is particularly suitable for 3D imaging of cleared tissues because of its 3D imaging ability and high imaging speed (Ahrens et al., 2013; Chen et al., 2014; Dodt, 2015; Dodt et al., 2007; Hillman et al., 2019; Huisken et al., 2004; Keller et al., 2010; Keller et al., 2008; Mano et al., 2018; Matsumoto et al., 2019; Planchon et al., 2011; Pernal et al., 2020; Power and Huisken, 2017; Tainaka et al., 2018; Tomer et al., 2014; Weber et al., 2014, Wu et al., 2011, Ueda et al., 2020). Essentially, the combination of tissue clearing techniques and LSM makes high-resolution 3D tissue imaging much more efficient and practical by replacing physical tissue sectioning with optical sectioning. Nevertheless, despite the potential benefits of the combination, it is still challenging to adopt them in 3D tissue imaging for several reasons.

First, conventional LSM is barely capable and inefficient to image centimeter-scale tissues with micron-scale or higher spatial resolution. In order to image a large sample with LSM at a micron-scale or higher spatial resolution, the sample is usually translated to multiple lateral positions to image a series of subdivided sample volumes in sequence (Figure 1a-b). The 3D image of the entire sample is obtained by registering and combining the collected 3D image stacks from all subvolumes. The strategy inherits the classic tradeoff between the spatial resolution and the size of the field of view (FOV) in LSM (Gao, 2015). The spatial resolution increases with the increase of the detection numerical aperture (NA) and the decrease of the excitation light sheet thickness, but the size of the detection FOV and the length of the excitation light sheet decrease at the same time (Figure 1a). In consequence, the sample is divided into more subvolumes to be imaged entirely (Gao et al., 2019). It not only increases the imaging acquisition time, but also makes the image registration and combination much more challenging, especially when the tissue is not rigid, which usually introduces structure distortions during imaging. Conversely, the size of the detection FOV and the length of the excitation light sheet must increase to decrease the number of subvolumes and the image acquisition time. However, the spatial resolution and optical sectioning ability drop at the same time (Figure 1b) (Tomer et al., 2014). Therefore, a major challenge of high-resolution 3D cleared tissue imaging using LSM is to achieve the desired spatial resolution while improving the image acquisition efficiency, reducing the data processing workload, and maintaining the image registration accuracy at the same time. Meanwhile spatial resolutions, that are suitable for imaging tissues of different sizes and required to answer biological questions at subcellular, cellular, and tissue levels, are often different. It is therefore necessary to be able to adjust the spatial resolution of a microscope conveniently to acquire spatial information at different scales efficiently. Conventional light sheet microscopes can hardly satisfy these requirements.

**Figure 1.**
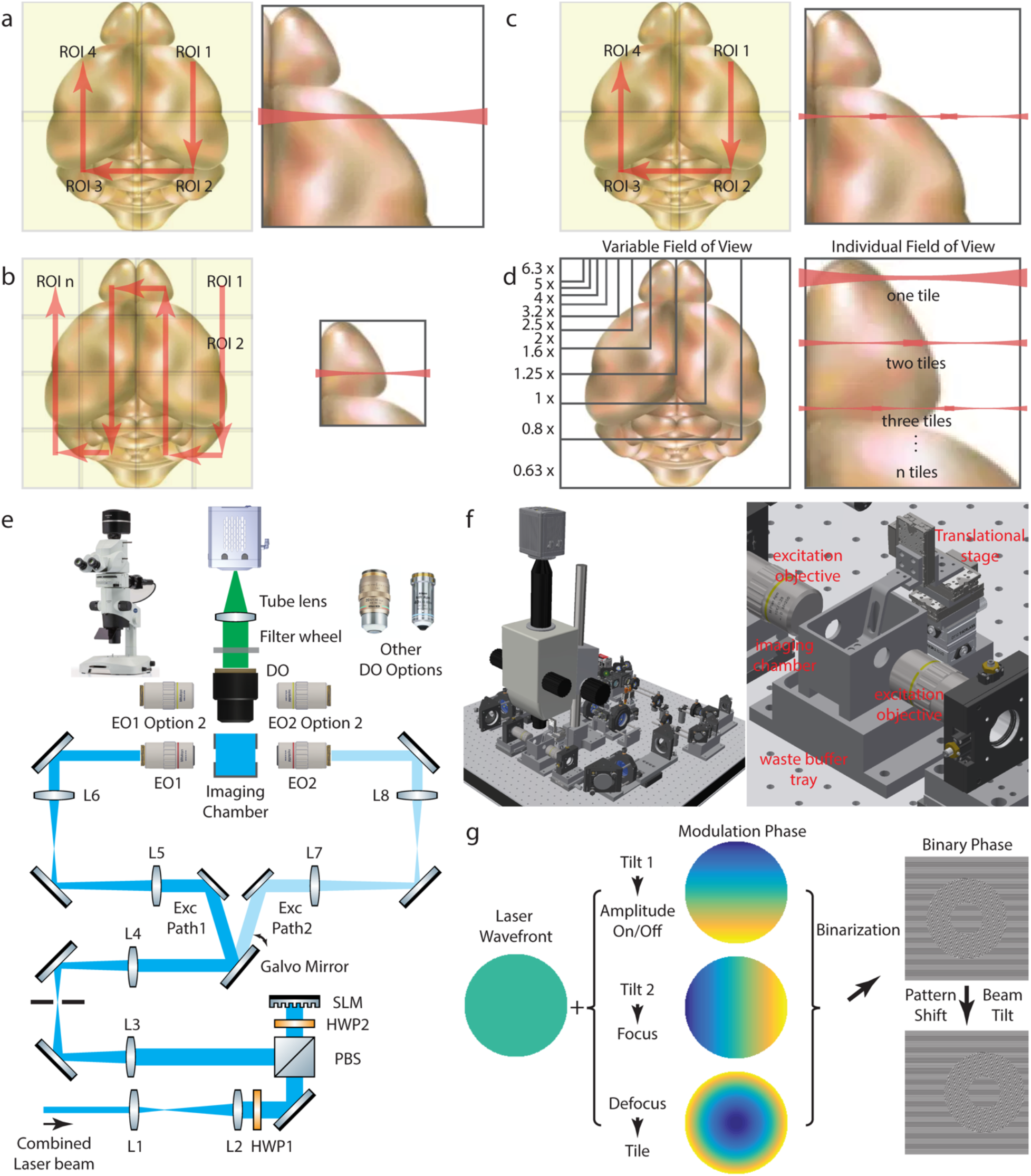
The tiling light sheet microscope for cleared tissue imaging. (a,b) Either the spatial resolution or the imaging speed is decreased when a large sample is imaged with conventional LSM. (c) Both the high spatial resolution and the high imaging speed are maintained by using tiling light sheets. (d) Both the size of the FOV and the excitation tiling light sheet can be adjusted based on the desired the spatial resolution and imaging speed. (e,f) The schematic diagram and configuration of the tiling light sheet microscope. (g) The generation of binary phase maps used to modulate the illumination light. Tilt 1 phase is used to control the excitation laser beam intensity profile. Tile 2 phase is adjusted to keep the excitation light sheet in focus within the FOV. Defocus phase is used to tile the excitation light sheet and to correct the light sheet lateral drifting caused by RI variation of the imaging buffer. All phase components are superimposed and binarized before being shifted to correct the tilt of the excitation beam.

Second, tissue clearing and the associated fixation and labeling techniques play essential roles in high-resolution cleared tissue imaging. Many tissue clearing methods have been developed in recent years (Aoyagi et al., 2015; Azaripour et al., 2016; Becker et al., 2012; Cai et al., 2019; Erturk et al., 2014; Gradinaru et al., 2018; Hahn et al., 2019; Hasegawa et al., 2016; Inoue et al., 2019; Jing et al., 2018; Ke et al., 2013; Ke et al., 2016; Tainaka et al., 2016; Ueda et al., 2020), but it is still difficult to make centimeter-scale tissues uniformly transparent. In addition, different types of biological tissues processed by different methods often preserve endogenous fluorophores differently, have different transparency, optical refractive indexes, mechanical strengths, hydrophilic properties and morphological distortions. These issues restrict the methods that can be used to clear and image specific tissues and limit the highest spatial resolution that can be obtained. Conventional light sheet microscopes are usually not compatible with all tissue clearing methods and can rarely be optimized to fit different tissue properties. These limitations not only prevent the use of the most suitable tissue clearing method in many applications but also hinder the development of new tissue clearing techniques. Thus, having the flexibility to image cleared tissues of different properties is essential to facilitate the development and application of tissue clearing techniques.

In addition to the problems above, it is necessary to lower the threshold to adopt tissue clearing and high-resolution 3D tissue imaging techniques in different applications. Unfortunately, most light sheet microscopes are either not optimized to image centimeter-scale tissues cleared by different methods with high spatial resolution, or too sophisticated to align and operate, regardless of the usually limited 3D imaging ability. Therefore, a reliable, flexible, easy to operate, and low-cost light sheet microscope with advanced 3D imaging ability is required to promote the adoption of tissue clearing and imaging techniques in different fields to unleash their potential.

Recently, multiple light sheet microscopes were developed specifically to image cleared tissues, but these problems still remain not fully addressed. For example, an open-top light sheet microscope was developed to improve the imaging throughput by simplifying the sample loading process (Glaser et al. 2019), but the tissue thickness is limited to a couple of millimeters and the spatial resolution is limited to a few microns. Axially scanned light sheet microscopes (ASLM) were developed to improve the spatial resolution (Chakraborty et al., 2019; Voigt et al., 2019). However, ASLM requires precise synchronization between the axially scanned light sheet and the rolling shutter of the detection camera, which restricts the use of the microscope only to uniformly transparent tissues. The synchronization requirement also makes the microscope very difficult to align and adjust. Microscope realignment is required to image cleared tissues with different RIs and to make any change in the spatial resolution, imaging speed, excitation light sheet geometry, and the size of the FOV. In addition, this technique suffers from severe photobleaching as the excitation light sheet becomes thinner and shorter to improve the spatial resolution. The use of the resonance galvanometer causes more photobleaching due to the extra sample illumination beyond the detection FOV. A light sheet theta microscope (LSTM) was developed to image cleared tissues more uniformly and to eliminate limits on lateral dimensions of the sample, but it has the same problems as that of ASLM due to the synchronization requirement between the scanning light sheet and the detection camera rolling shutter (Migliori et al., 2018).

Here we present a novel tiling light sheet microscope to address the above problems. Different from other light sheet microscopes, the design of our microscope not only focuses on its 3D imaging ability to acquire the desired result from large cleared tissues, but also on its capability to optimize and adjust the imaging ability in a wide range, so that the microscope can be used conveniently in various applications. New optical design and phase modulation methods are used in the microscope to achieve a more advanced 3D imaging ability and reliability for cleared tissue imaging than the tiling light sheet microscope reported previously (Gao, 2015; Fu et al., 2016). The microscope has several unique features compared to the existing light sheet microscopes. First, tiling light sheets are used for sample illumination, so that the spatial resolution and imaging efficiency are improved at the same time compared to conventional LSM (Figure 1c). Second, the detection NA, detection magnification and the excitation light sheet can be adjusted easily, so that the microscope is capable to image centimeter-scale cleared tissues with micron-scale to nanometer-scale spatial resolution (Figure 1d). Third, the illumination light is phase modulated by a binary spatial light modulator (SLM) to create and optimize tiling light sheets for sample illumination. Moreover, the phase modulation through the SLM is used to correct almost all alignment errors and enables semi-automatic alignment of our microscope, which makes the microscope reliable and easy to operate. Finally, the microscope is compatible with all tissue clearing methods and capable to image centimeter-scale tissues of different shapes and mechanical strengths. Altogether, we deliver a tiling light sheet microscope for cleared tissue imaging. We demonstrate its advanced multicolor 3D imaging ability by imaging various tissues cleared by different methods with micron-scale to nanometer-scale spatial resolution.

## Results

### Microscope design and working principle

The configuration of the microscope is shown in Figure 1e-f. A binary SLM conjugated to a galvanometer is used to modulate the illumination light. The galvanometer directs the illumination light to one of the two symmetrical illumination paths and scans the laser beam to create a virtual excitation light sheet for sample illumination. The modulated laser beam is further conjugated to the rear pupils of both excitation objectives and illuminate the sample from one of the two opposite directions to minimize the illumination light travel distance in the sample to reach the detection FOV. Air objectives are used to illuminate the sample and to avoid contamination by the imaging buffer. The emission fluorescence is collected by a switchable detection objective and imaged onto a sCMOS camera through an optical filter. The detection magnification is adjustable from 0.63× to 6.3× to change the size of the detection FOV. The sample is immersed in RI matched imaging buffer and driven by a 3D translational stage for 3D imaging.

The microscope has four imaging modes (Figure 1c-d, figure supplement 1). (1) 2D imaging of a selected sample plane. (2) 3D imaging of a selected sample volume, in which a subvolume is imaged with tiling light sheets by scanning the sample in the axial direction. (3) 3D imaging of multiple selected sample volumes. (4) 3D imaging of an array of sample volumes. The microscope needs to be calibrated via software based on the imaging buffer RI before imaging (Figure 1g, figure supplement 2). A group of phase maps is generated next according to the desired spatial resolution and imaging speed by combining multiple phase components that are used to control the excitation beam intensity profile, tile the light sheet, focus the light sheet and correct alignment errors respectively (figure supplement 3). The generated phase maps are applied to the SLM in sequence repetitively and synchronized with camera exposures during imaging (figure supplement 1).

### 3D tissue imaging with micron-scale spatial resolution

Our tiling light sheet microscope uses low NA, long working distance (WD) air detection objectives to image cleared tissues with micron-scale spatial resolution. It not only makes the microscope compatible with different tissue clearing methods without being contaminated by the imaging buffer but also reduces the microscope cost and maintenance requirement. On the other hand, nearly isotropic micron-scale spatial resolution can still be achieved by using thin tiling light sheets despite the low detection NA, which is very difficult for conventional LSM (figure supplement 4,5). We evaluated the imaging ability of the microscope to image with micron-scale spatial resolution and compared it with a conventional light sheet microscope by imaging a Thy1-YFP mouse brain cleared with PEGASOS (Figure 2, Video 1) (Feng et al., 2000; Jing et al., 2018). A conventional light sheet microscope (Zeiss Z1) equipped with a 0.16 NA air objective was first used to image the sample at ~2×2×15 μm^3^ spatial resolution in ~1.5 hours. Next, the same mouse brain was imaged using our microscope equipped with a 0.25 NA, 60 mm WD air objective in both a six-tile mode at ~2×2×5 μm^3^ spatial resolution in ~5 hours and a non-tile mode at ~4×4×10 μm^3^ spatial resolution in ~12 mins. A ~5 μm thick light sheet was tiled six times in a ~2×2 mm^2^ FOV in the six-tile mode, and a ~10 μm thick light sheet was used to illuminate a ~4×4 mm^2^ FOV without tiling in the non-tile mode. The cleared mouse brain was imaged with either higher spatial resolution or faster imaging speed using our microscope. It is worth mentioning that it would take ~27 hours to image the mouse brain with the Z1 microscope at the same spatial resolution as that of the six-tile mode of our microscope because a thinner and shorter light sheet must be used to illuminate the sample, which increases the number of subvolumes by at least 6 times and collects ~18 times more images. Even the axial resolution of the result collected in the non-tile mode of our microscope is higher than that of the result collected with the Z1 microscope. The reason is that the lateral position of the excitation light sheet drifts when the RI of the imaging buffer varies to image tissues cleared by different methods. The lateral position of the excitation light sheet must be adjusted to center the light sheet in the FOV so that the shortest and thinnest light sheet can be used to illuminate the sample. Otherwise, a longer but thicker light sheet must be used to keep the spatial resolution uniform within the FOV for the light sheet microscopes without the ability to correct the light sheet lateral drift, which then inevitably reduces the axial resolution (figure supplement 6). Light sheet lateral drifting is also a major problem that makes it practically difficult to image cleared tissues at high spatial resolution with conventional LSM despite its possibility in theory because it is more challenging to compensate the light sheet drift as the light sheet becomes thinner and shorter.

**Figure 2.**
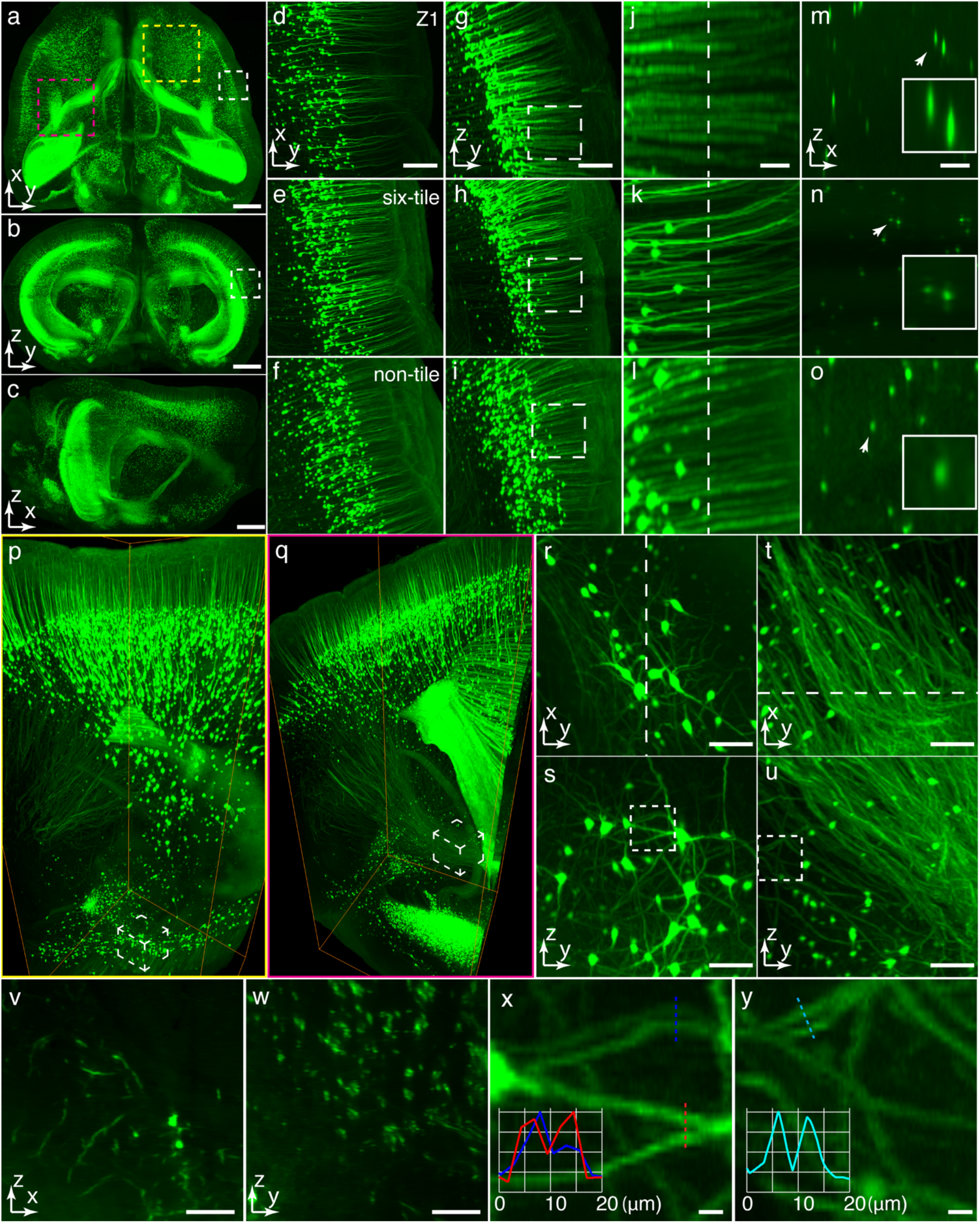
The 3D tissue imaging ability evaluation and comparison with a conventional light sheet microscope. (a-c) Lateral and axial maximum intensity projections (MIP) of a cleared Thy1-YFP mouse brain imaged with the presented microscope. (d-f) Lateral and (g-i) axial MIPs of a selected volume (white) in (a) and (b), imaged with a Zeiss Z1 light sheet microscope, a six-tile mode and a non-tile mode of the presented tiling light sheet microscope. (j-l) Zoom in views of the selected areas in (g-i). (m-o) Axial slices through the indicated planes in (j-l). (p,q) 3D renderings of two ~2×2×5 mm^3^ subvolumes (yellow and magenta) in (a) imaged with a six-tile mode of the microscope. (r-u) Lateral and axial MIPs of the selected volumes in (p) and (q). (v,w) Axial slices through the indicated planes in (r) and (t). (x,y) Zoom in views of the selected areas in (s) and (u). The size of all inserts in (m-o) is 50×50 μm^2^. Scale bars, 1 mm (b-d), 200 μm (d,g), 50 μm (j,m), 100 μm (r-w), and 10 μm (x,y).

3D tissue imaging with the nearly isotropic micron-scale spatial resolution is well suitable for visualizing tissue structures at the cellular level. For instance, the vascular network of a mouse brain (SMACre::Ai14 cleared with PEGASOS) with nearly every capillary recognizable was acquired in ~3 hrs (Video 2). It is very difficult to obtain comparable results with conventional LSM as shown in the comparison (Figure 3a-i). The result collected with our microscope provides a clear topological understanding of the brain vascular architecture. It also shows that the penetrating vessels in the cerebral cortex connect to venules or arterioles in the hippocampus at the corpus callosum via capillaries (Figure 3j-q, Video 2), which wasn’t directly observed before. The advanced 3D imaging ability of our microscope could facilitate the understanding of brain-wide vascular networks crossing distinct brain regions. In addition to the advanced 3D imaging ability, various sample mounting methods were developed to image tissues of different shapes, sizes, and mechanical strengths (Figure 3r-w, figure supplement 7-13), which allows the use of the microscope in various studies.

**Figure 3.**
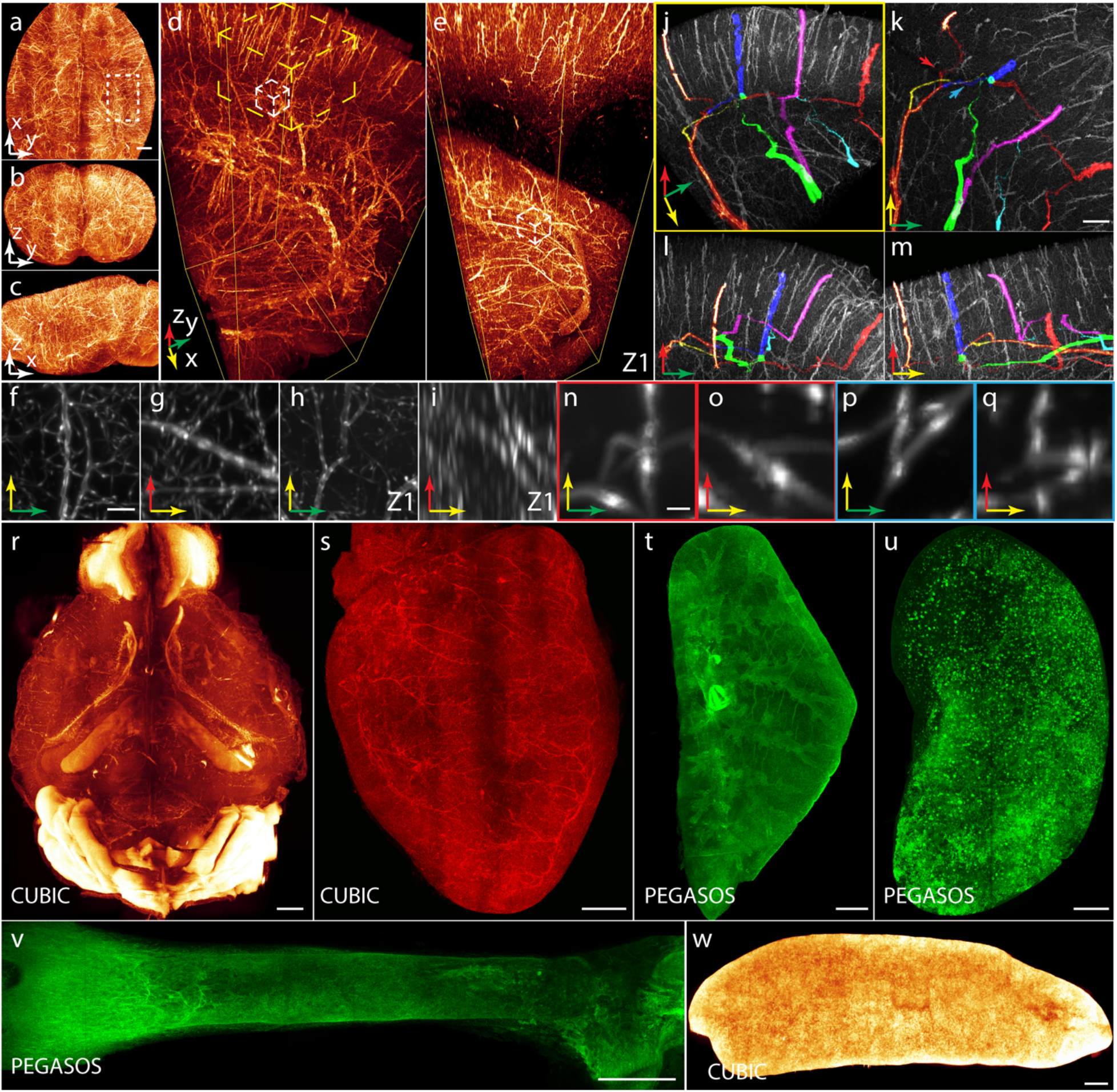
3D imaging of mouse organs cleared by different methods with micron-scale spatial resolution. (a-c) Lateral and axial MIPs of a cleared mouse brain imaged with the microscope showing the mouse brain vascular network. (d) 3D rendering of the selected subvolume in (a). (e) 3D rendering of a similar volume as that in (a) imaged with a Zeiss Z1 microscope. (f,g) Lateral and axial MIPs of the selected volume (white) in (d). (h,i) Lateral and axial MIPs of the selected volume in (e). (j) 3D rendering of the selected volume (yellow) in (d) with colored traces showing connected penetrating vessels and hippocampus vessels at corpus callosum via capillaries. (k-m) Lateral and axial MIPs of the volume in (j). (n-q) Lateral and axial MIPs of the regions indicated by arrows in (k), showing the zoom in views of the connecting capillaries. (r-w) 3D renderings of various mouse organs cleared with CUBIC and PEGASOS, including mouse brain, heart, lung, kidney, femur and spleen. Scale bars, 1 mm (a, r-w), 40 μm (f), 200 μm (k) and 10 μm (n).

### Multicolor 3D tissue imaging

Multicolor 3D imaging is required to determine the relative distribution of different organelles in tissues. Multicolor 3D imaging is usually realized by the sequential imaging of the same sample volume with illumination lasers of different wavelengths and emission filters. The collinear alignment of different lasers beams is critical to ensure an accurate image registration of different color channels. It is ensured by phase modulation of the illumination light at all wavelengths in our microscope (Figure 1g, figure supplement 3). We examined the multicolor 3D imaging ability of the microscope by imaging a dual color labeled mouse mammary gland. The sample was first immune-stained with Krt14/AF488, a mammary basal cell marker, and Cd200/AF594, expressed in endothelial, stromal, and part of the luminal cells, then it was cleared with Adipo-clear (Chi et al., 2018). The results show that Cd200 stained luminal cells localize on the inner layer of the mammary ducts, the outer layer of which is marked by Krt14 stained basal cells. (Figure 4a-s, Video 3). The relative expression of Krt14, labeling stain basal cells, and Cd200, labeling stromal and blood vessels, also shows the distinct patterns at mammary terminal end buds and ducts (Figure 4t-y), this suggests the spatial heterogeneity of the mammary cell composition. In another example, we imaged a human breast cancer tissue and an adjacent normal breast tissue. Both samples were immuno-stained with Krt14/AF488 and Cd146/AF594 and cleared with Adipo-clear (Figure 4z-gg). The cancer tissue lost its alveoli with abnormal tube structures and shows a denser vascular system that likely favors the intravasation of cancer cells. The patient had enlarged lymph nodes with suspicious metastatic cells, which is consistent with the abnormal vasculature pattern. The experiment suggests the prognostic value of tissue clearing and high-resolution 3D tissue imaging techniques in pathological applications.

**Figure 4.**
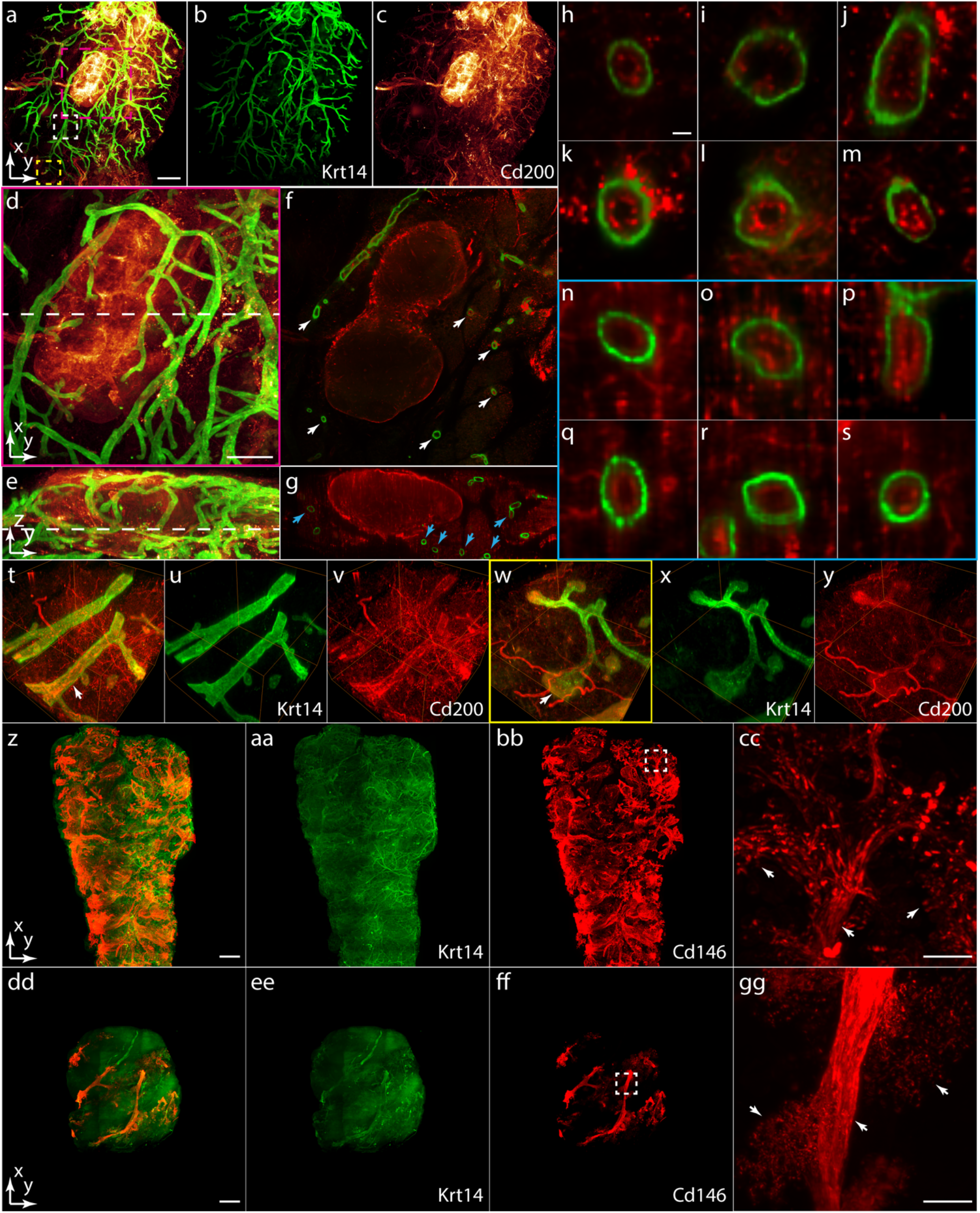
Multicolor 3D cleared tissue imaging. (a-c) 3D renderings of a cleared dual color immuno-stained mouse mammary gland, rendered in both colors together and individually. (d,e) Lateral and axial MIPs of the selected volume (magenta) in (a). (f,g) Lateral and axial slices through the indicated planes in (e) and (d). (h-s) Zoom in views of the lateral and axial cross sections of mammary ducts indicated in (f) and (g) showing the relative distributions of Krt14 and Cd200 labeled cells. (t-y) 3D renderings of two selected volumes (white and yellow) in (a), showing distinct patterns of Krt14 and Cd200 at mammary ducts and end buds. (z-bb) 3D renderings of a cleared dual color immuno-stained human breast cancer tissue. (cc) Zoom in view of the selected area in (bb). (dd-ff) 3D renderings of a cleared dual color immuno-stained human breast nomal tissue adjacent to cancer tissue in (z). (gg) Zoom in view of the selected area in (ff). Scale bars, 1 mm (a,z,dd), 500 μm (d), 20 μm (h), 200 μm (cc,gg).

### 3D tissue imaging with submicron scale spatial resolution

In order to visualize subcellular tissue structures, the spatial resolution of the microscope can be improved to submicron scale by using high-NA detection objectives and thinner tiling light sheets. Both illumination objectives and the detection objective of the microscope are replaceable, so that high-NA objectives, up to 1.0 NA, and thinner tiling light sheets, down to ~1 μm light sheet thickness, can be used to improve the 3D imaging ability of the microscope without realigning the microscope. Only a new group of phase maps need to be generated to operate the microscope under new imaging conditions. We verified the improved 3D imaging ability by imaging two Thy1-eGFP mouse brains, cleared with PEGASOS and CUBIC respectively (Figure 5, Video 4) (Jing et al., 2018; Susaki et al., 2014; Susaki et al., 2015). A 1.0 NA, 8.2 mm WD immerse objective with RI correction ability from 1.44 to 1.50 was used to collect the emission fluorescence. We imaged two similar regions in both samples at ~0.3×0.3×1.5 μm^3^ spatial resolution with a ~2.5 μm thick tiling light sheet tiled at four positions in a ~300×300 μm^2^ FOV. As expected, subcellular neuronal structures were much better resolved than the result obtained with the 0.25 NA air objective, indicated by clearly resolved dendritic spines on the excitatory pyramidal neurons, neuron axons and the morphology of individual neurons. However, both the image acquisition time and data size increased proportionally to the 3D spatial resolution. The spatial resolution improvement from ~2×2×5 μm^3^ to ~0.3×0.3×1.5 μm^3^ results in roughly a one hundred time increase in image acquisition time and data size for the same sample volume. Therefore, the spatial resolution in cleared tissue imaging should be determined according to the required spatial resolution and sample size despite the imaging ability of the microscope. Appropriate spatial resolution can be set conveniently by changing the detection NA, magnification and the excitation tiling light sheet with our tiling light sheet microscope.

**Figure 5.**
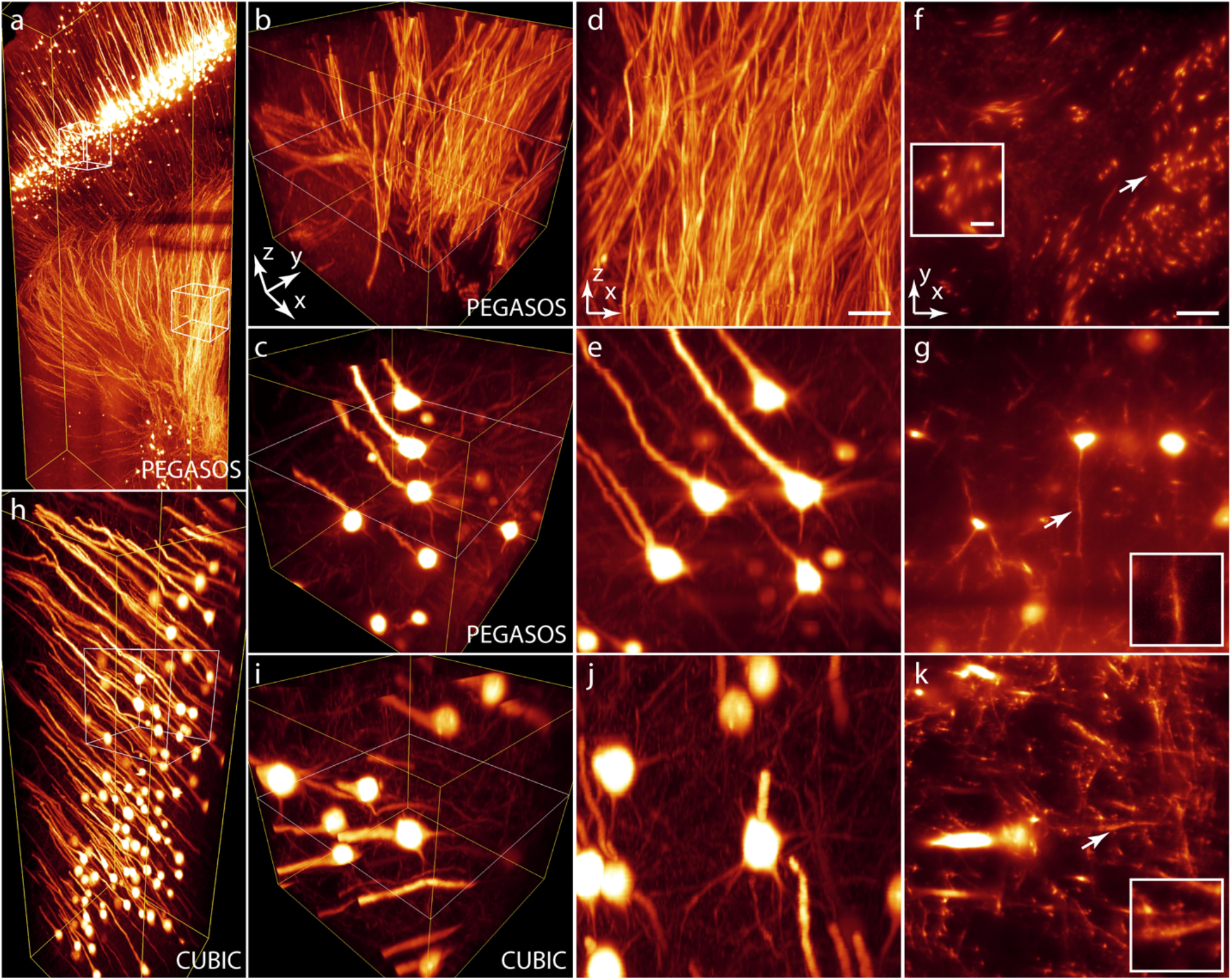
3D tissue imaging with submicron spatial resolution and comparison of different tissue clearing methods. (a) 3D rendering of a ~0.3×1.1×1.5 mm^3^ sample volume in a Thy1-eGFP mouse brain cleared with PEGASOS, which was reconstructed from four ~0.3×0.3×1.5 mm^3^ subvolumes. (b,c) 3D renderings of two selected ~0.15×0.15×0.15 mm^3^ subvolumes in (a). (d,e) Axial MIPs of the subvolumes in (b) and (c). (f,g) Lateral slices through the indicated planes in (b) and (c). (h) 3D rendering of a ~0.3×0.3×1.5 mm^3^ sample volume in a Thy1-eGFP mouse brain cleared with CUBIC. (i) 3D rendering of a selected 0.15×0.15×0.15 mm^3^ subvolume in (h). (j) Axial MIP of the subvolume in (i). (k) Lateral slice through the indicated planes in (i). The size of all inserts in (f,g,k) is 20×20 μm^2^. Scale bars, 20 μm (d, f), 5 μm (f) insert.

The results also reveal the differences between these tissue clearing methods. The mouse brain cleared with CUBIC was expanded by ~16% in each dimension compared to its original size. The tissue autofluorescence was lower and the endogenous fluorescence was better preserved compared to the sample cleared with PEGASOS. Thus, neuronal structures were better resolved as indicated by the higher image contrast and the more clearly visualized dendritic spines (Figure 5e-k, Video 4). However, tissues cleared with CUBIC are usually very soft and the CUBIC imaging buffer has a relatively high viscosity, so that the sample could deform during sample translations. The distortions make it extremely difficult to register the 3D images of different subdivided sample volumes at submicron spatial resolution. On the contrary, the mouse brain cleared with PEGASOS shrank by ~25% in each dimension, but the sample is rigid enough to prevent distortions, which enables accurate image registration and combination at submicron spatial resolution, despite the heavier fluorescence quenching and stronger autofluorescence background from our observation. Fortunately, our tiling light sheet microscope is compatible with both tissue clearing methods, so that the most appropriate tissue clearing method can be used to clear the sample according to need of the research.

### 3D tissue imaging with submicron spatial resolution via tissue expansion

Tissue expansion techniques provide a solution to resolve tissue structures with a higher spatial resolution than that of the microscope (Chang et al., 2017; Chen et al., 2015; Gao et al., 2019; Ku et al., 2016; Tillbery et al., 2016; Wassie et al., 2019). The equivalent spatial resolution corresponding to the resolving ability improves proportionally to the tissue expansion ratio. We investigated the imaging ability of the presented tiling light sheet microscope in combined with tissue expansion techniques. A Thy1-eGFP mouse brain was cleared and expanded by ~5 times in each dimension using MAP (figure supplement 14) (Ku et al., 2016). A ~9×11×5 mm^3^ volume in the hippocampus of the expanded mouse brain was imaged with the 0.25 NA air objective at ~2×2×5 μm^3^ spatial resolution using water as the imaging buffer. The equivalent spatial resolution corresponding to the resolving ability is ~0.4×0.4×1 μm^3^ due to the ~5 times expansion ratio. As shown (Figure 6, Video 5), both the cellular and subcellular neuronal structures in the hippocampus of the mouse brain are well visualized, indicated by the clearly resolved individual neuronal axons and dendritic spines despite the high cell density in the hippocampus. Dashed line pattern can be observed from some neuron axons, which is likely caused by axoplasmic transport in these neurons. The expanded mouse brain is also sufficiently rigid to avoid deformations caused by sample translation during imaging, which enables accurate image registration of the subdivided sample volumes. Convincingly, the combination of our microscope and tissue expansion techniques offers an alternative solution to study cell morphology and organization in large samples with submicron spatial resolution, such as the axonal and dendritic morphology of neurons and long-distance neuron axonal projections across multiple brain regions.

**Figure 6.**
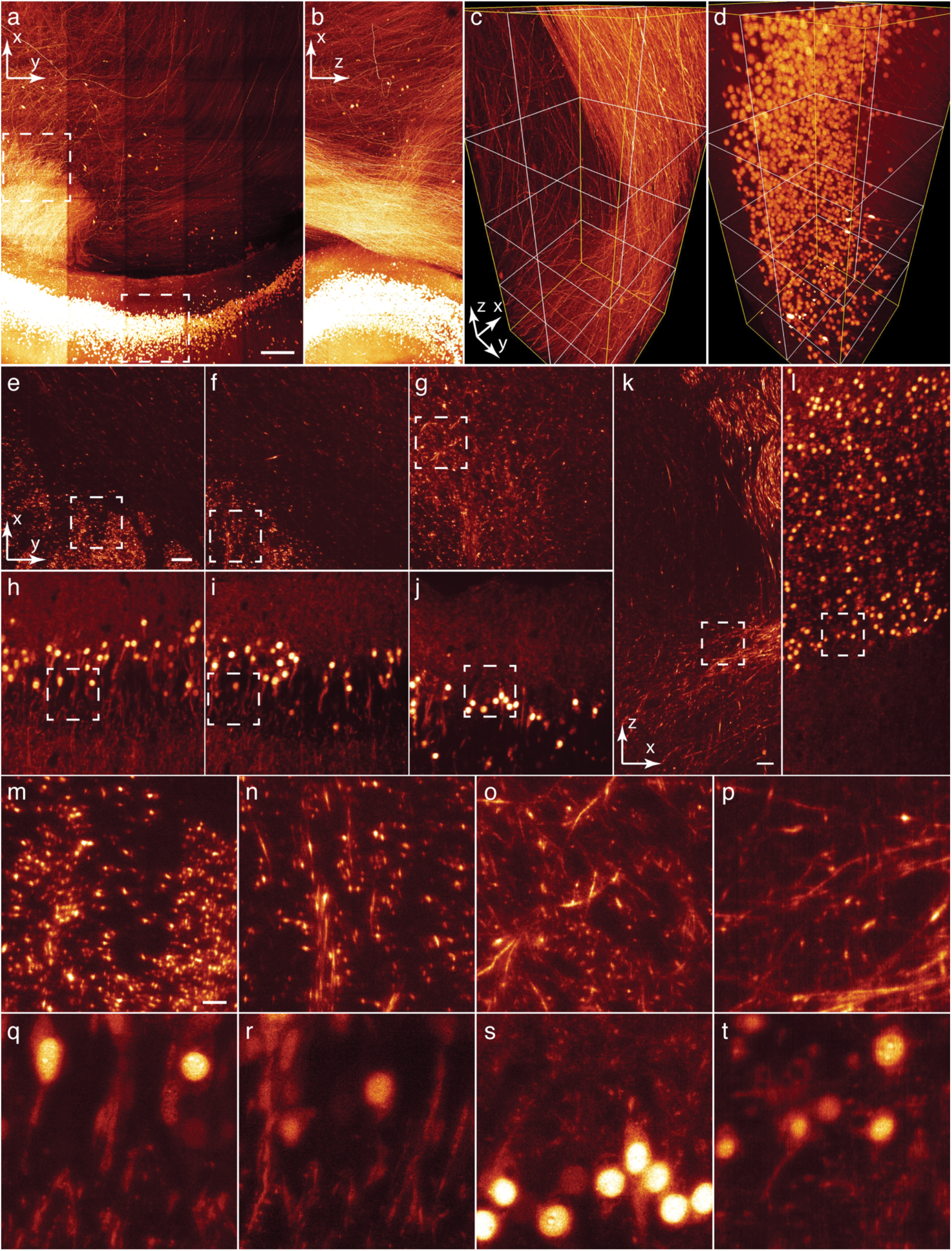
3D tissue imaging with submicron spatial resolution via tissue expansion. (a) 3D rendering of a ~9×11×5 mm^3^ sample volume in a ~5 times expanded Thy1-eGFP mouse brain, reconstructed from thirty ~2×2×5 mm^3^ subvolumes. (b) Axial MIP of the sample volume in (a). (c,d) 3D renderings of two selected ~2×2×5 mm^3^ subvolumes in (a). (e-g) Lateral slices through the indicated lateral planes in (c) from top to bottom. (h-j) Lateral slices through the indicated lateral planes in (d) from the top to the bottom. (k,l) Axial slices through the indicated axial planes in (c) and (d). (m-p) Zoom in views of the selected areas in (e-g and k). (q-t) Zoom in views of the selected areas in (h-j and l). Scale bars, 1 mm (a), 200 μm (e, k), 50 μm (m).

### 3D tissue imaging with nanometer-scale spatial resolution

The microscope is capable to further resolve tissue structures with nanometer-scale spatial resolution by imaging expanded tissues with high-NA detection objectives and thinner tiling light sheets. To demonstrate the ability, we first imaged a ~2.3×2.3×2 mm^3^ sample volume in an expanded Thy1-eGFP mouse brain at ~2×2×5 μm^3^ spatial resolution with the 0.25 NA air objective. Next, we replaced the air objective with a 0.8 NA, 3.5 mm WD water immerse objective and imaged a ~0.3×0.3×0.6 mm^3^ region within the imaged volume at ~0.33×0.33×1 μm^3^ spatial resolution. The spatial resolutions corresponding to the resolving abilities are ~0.4×0.4 ×1 μm^3^ and ~70×70×200 nm^3^ respectively due to the ~5 times tissue expansion ratio. As shown (Figure 7a-j, Video 6), the result obtained with the 0.8 NA water immerse objective reveals significantly more neural structures because of the higher detection NA and thinner excitation tiling light sheets, which improve the spatial resolution and the optical sectioning ability at the same time. The results suggest that brain neural networks are extremely complicated. 3D imaging at nanometer-scale spatial resolution might be necessary to resolve the brain neural networks completely. It could be extremely challenging due to the problems of photobleaching, imaging acquisition time, image processing workload and data management. Fortunately, our microscope enables the tuning of the imaging ability in a wide range to image the sample with necessary spatial resolution while minimizing the associated problems.

**Figure 7.**
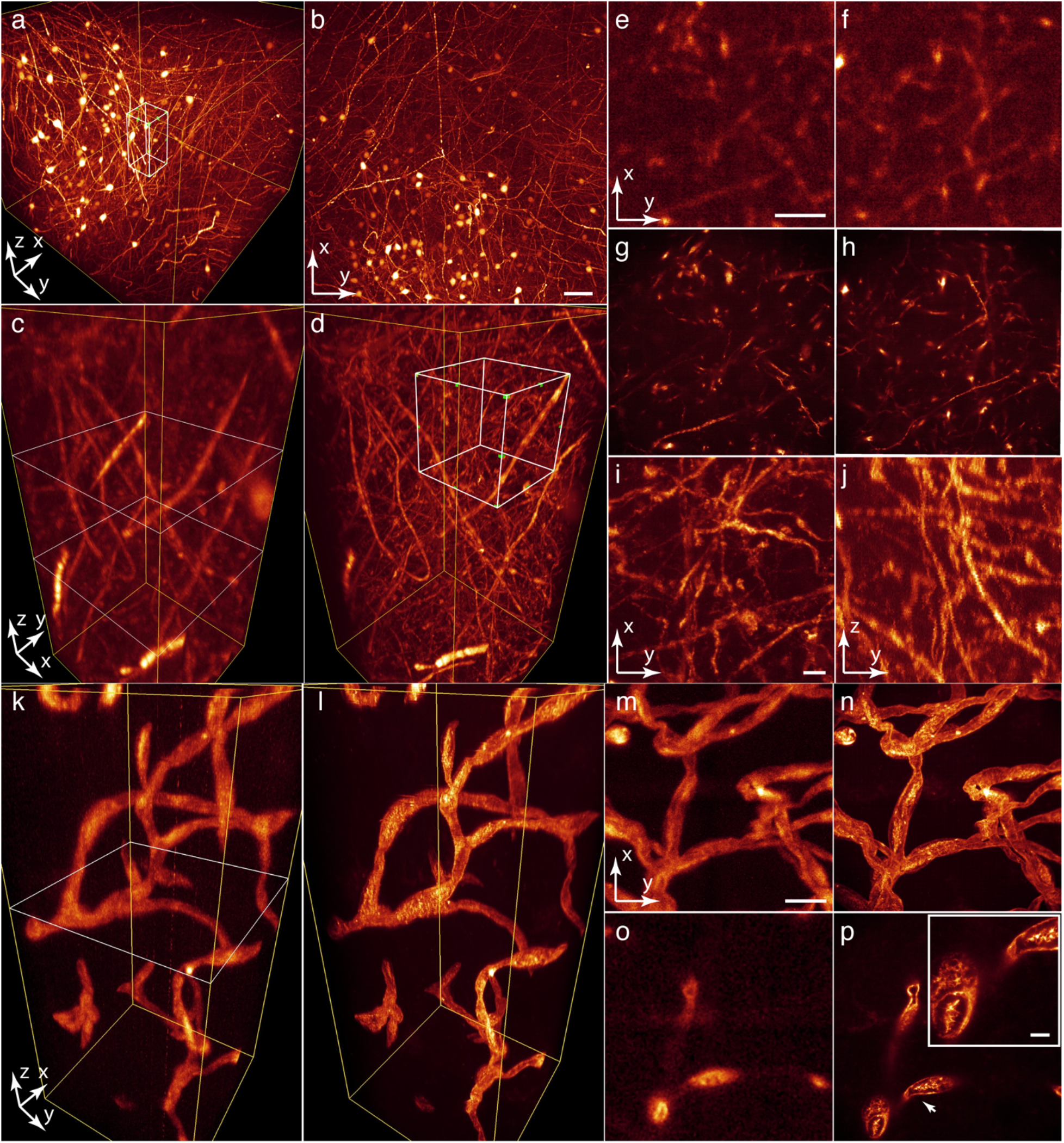
3D tissue imaging with nanometer-scale spatial resolution. (a) 3D rendering of a ~2.3×2.3×2 mm^3^ sample volume in a ~5 times expanded Thy1-eGFP mouse brain imaged with a submicron spatial resolution. (b) Lateral MIP of the sample volume in (a). (c) 3D rendering of the selected ~250×250×500 μm^3^ subvolume in (a). (d) 3D rendering of the same subvolume in (c) imaged with a nanometer-scale spatial resolution. (e,f) Lateral slices through the indicated lateral planes in (c) from top to bottom. (g,h) Lateral slices through the same planes as (e) and (f) in (d). (I,j) Lateral and axial MIPs of the selected sample volume in (d). (k,l) 3D renderings of the same ~250×250×600 μm^3^ sample volume in a ~5 times expanded CAG-zsGreen mouse brain imaged with a submicron spatial resolution and a nanometer-scale spatial resolution. (m,n) Lateral MIPs of the sample volume in (k) and (l). (o) Lateral slice through the indicated lateral plane in (k). (p) Lateral slices through the same lateral planes as (o) in (l). Scale bars, 200 μm (b), 50 μm (e,m) and 10 μm (i,p insert).

Discontinuous light sheets created by scanning coaxial beam arrays synchronized with the detection camera rolling shutter can be used to improve the imaging speed of the microscope without decreasing the spatial resolution (Wang et al., 2019). In order to keep the scanning coaxial beam array and the detection camera rolling shutter in sync, the sample must be uniformly transparent. Expanded tissues are sufficiently uniform and transparent to be imaged with the discontinuous light sheets. A ~3.8×3.8×7 mm^3^ sample volume in an expanded CAG-zsGreen mouse brain was imaged with the 0.25 air objective at ~2×2×4 μm^3^ spatial resolution in ~28 mins (Video 7). A three-waist discontinuous light sheet was tiled three times at each image plane to image the sample. It is three times faster than that using a regular tiling light sheet which would require nine tiles per image plane. Furthermore, both the vascular network in the mouse brain and the detailed structure of blood vessels can be visualized using our microscope with appropriate spatial resolutions from a submicron scale to a nanometer-scale (Figure 7k-p, Video 7).

### 3D tissue imaging with multiple resolution scales

The advantages of our tiling light sheet microscope could benefit researches using other research organisms. Capable of rapid whole-body regeneration from a tiny piece, the planarians Schmidtea mediterranea have become model animals to study stem cell functions and molecular mechanisms in tissue regeneration (Elliott et al.,2013; Reddien et al.,2018; Forsthoefel et al. 2020). In order to study the distributions and cell-cell interactions of stem cells in planarians, sophisticated 3D tissue imaging techniques are required due to the large body size and opaqueness of planarians. Both conventional LSM and tissue expansion techniques have been used individually in planarian studies (Wang et al.,2016; Davies et al.,2017; Lim et al.,2019), but the potential of these techniques have not been fully exploited. Our microscope provides a unique capability to image cleared tissues at multiple resolution scales in one microscope. A planarian co-stained with smedwi-1, a pan-neoblast marker labeling stem cells (Reddien et al.,2005), and propidium iodide (PI), a nuclear dye labeling all cell nuclei, was cleared, expanded and imaged with our microscope. At first, the stem cell distribution in the planarian was obtained by imaging the ~5 times expanded planarian in two colors with the 0.25 NA air objective at ~2×2×5 μm^3^ spatial resolution in ~3 hours (Figure 8). Next, we imaged a selected 1 mm^3^ region in the expanded planarian with the 0.8 NA water immerse objective to visualize the organizations and cell-cell interactions of the stem cells at ~0.33×0.33×1 μm^3^ spatial resolution in another four hours. As shown, the cell distribution regarding the smedwi-1+ labeled stem cells and the cell types surrounding a single smedwi-1+ cell can be clearly visualized, which are of interest but have not been fully addressed for years. Apparently, the distributions and functions of stem cells in planarians can be better studied with our tiling light sheet microscope. Similar advantages may apply to other research organisms, such as *C. elegans*, Drosophila melanogaster, zebrafish, water bear *Tardigrades*, *Macrostomum lignano*, and *Hofstenia miamia* (Yu et al. 2020).

**Figure 8.**
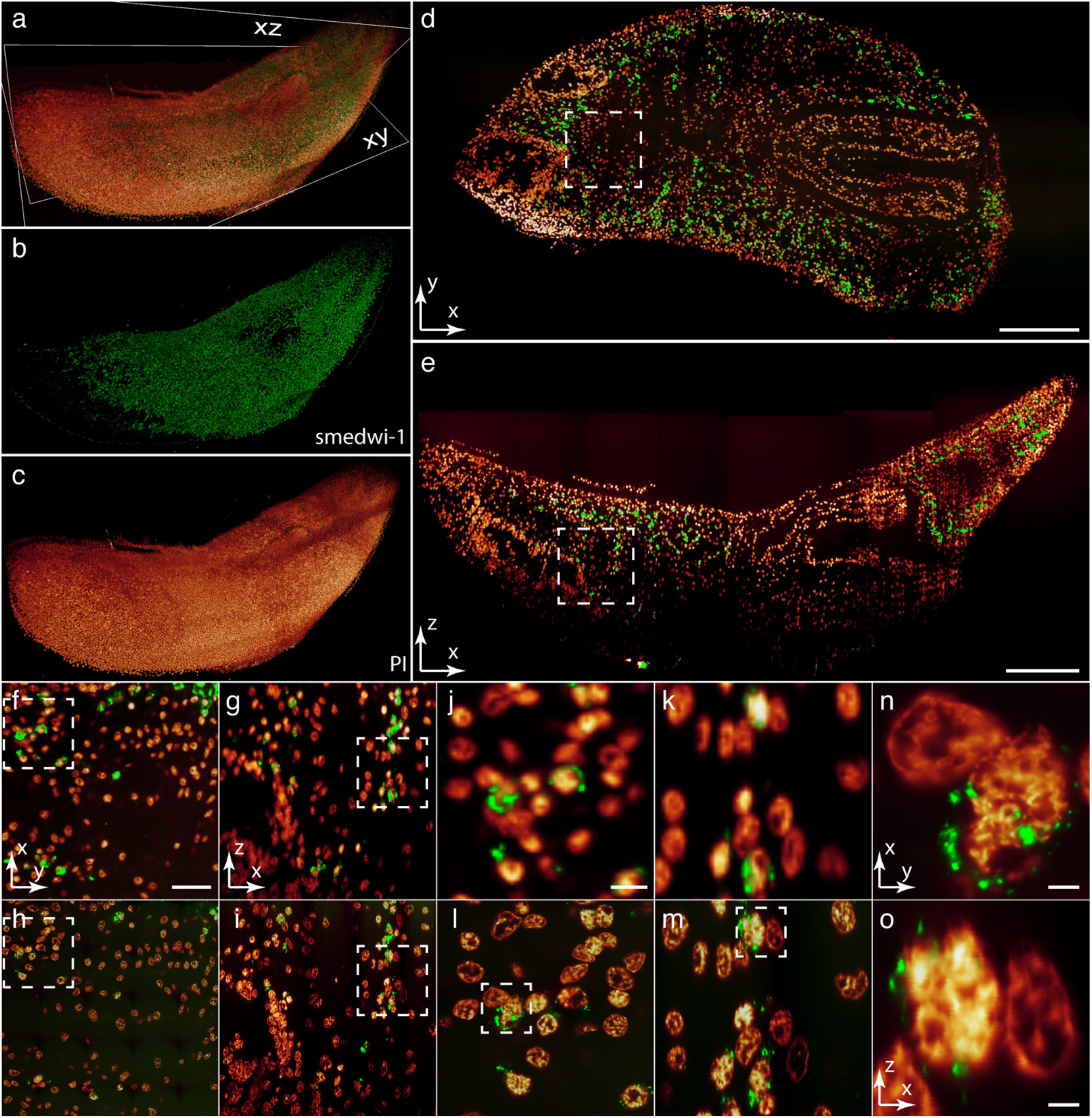
The stem cell distribution and organization in a planarian Schmidtea mediterranea. (a-c) 3D renderings of a ~5 times expanded planarian, labeling stem cells and cell nuclei, imaged with a submicron spatial resolution, rendered in two colors together and individually. (d,e) Lateral and axial slices through the indicated planes in (a). (f,g) Zoom in views of the selected areas in (d) and (e). (h,i) Lateral and axial slices through the same planes as (f) and (g) from the 3D image of the selected 1 mm^3^ volume imaged with a nanometer-scale spatial resoluton. (j-m) Zoom in views of the selected areas in (f-i). (n,o) Zoom in views of the selected areas in (l) and (m). Scale bars, 1 mm (d,e), 200 μm (f), 50 μm (j) and 10 μm (n).

## Discussion

As shown, our tiling light sheet microscope is capable of imaging centimeter-scale cleared tissues rapidly with micron-scale to nanometer-scale spatial resolution. While using real-time optimized tiling light sheets, nearly isotropic micron-scale and submicron scale spatial resolutions can be obtained with low-NA, long-WD air objectives and high-NA immerse objectives. The resolving ability can be further improved to a nanometer-scale by adopting tissue expansion techniques. In addition, the microscope can be adjusted conveniently to adapt to different requirements without realignment, which offers a unique ability to observe and correlate tissue structures at multiple resolution scales.

Although submicron spatial resolution can be achieved with our microscope by either using high-NA detection objectives and thinner light sheets or by adopting tissue expansion techniques, each approach has its advantages and disadvantages. First, tissue transparency and RI consistency are more critical in order to use high-NA objectives, because the performances of high-NA objectives are more easily affected by optical aberrations and light scattering. Unfortunately, it is still challenging to make tissues of centimeter-scales uniformly transparent in our experiences. Optimization of the tissue clearing method in specific applications is usually necessary, and more advanced tissue clearing methods are expected to address the issue. On the other hand, expanded tissues are more transparent and introduce very little optical aberrations. Second, high-NA, long-WD immerse objectives with RI correction ability above 1.4 are rare and expensive. RI correction according to the tissue to be imaged and the imaging buffer is always required before imaging to achieve the desired spatial resolution. On the contrary, low-NA, long-WD air objectives used to image expanded tissues are more available and affordable. RI correction is not required because water is always used as the imaging buffer and low detection NA is less vulnerable to the optical aberrations introduced by the air-water interface (figure supplement 6). Third, the lateral resolution increases faster than the axial resolution as the detection NA increases, which makes it more difficult to keep the spatial resolution isotropic. Differently, tissue expansion improves the resolving ability in both the lateral and axial directions by the same amount, which equals the tissue expansion ratio. Therefore, 3D tissue imaging with submicron spatial resolution is more feasible by adopting tissue expansion techniques. Nevertheless, tissue expansion requires more procedures and longer time to have the sample processed, which usually causes more losses of endogenous fluorophores. On the other hand, Low-NA objectives collect less fluorescence than high-NA objectives due to the smaller NA. Thus, the brightness and labeling density of the endogenous fluorophores are the key factors to determine whether tissue expansion can be used to improve the resolving ability in 3D tissue imaging.

Despite the imaging ability of the microscope, the spatial resolution should be determined wisely in specific applications. Tissue fixation and clearing introduce morphological distortions (Hama et al., 2015; Park et al., 2018). The reliability of the resolvable tissue structures with the selected spatial resolution should be validated carefully. The image acquisition time and the amount of data increase proportionally with the 3D spatial resolution and tissue size. It is typical to spend a few hours and collect hundreds of gigabytes to terabytes of data to image centimeter-scale samples at cellular spatial resolution. It comes as no surprise that petabytes of data must be collected to image centimeter-scale samples with submicron-scale or higher spatial resolutions. Data management and image processing capacities should be considered prior to imaging.

Photobleaching becomes an issue as the spatial resolution improves and the tissue size increases. There are two reasons. One is that the number of fluorophores in each voxel decreases as the spatial resolution improves. The other is the much larger sample size compared to the length of the light sheet waist (figure supplement 15). In LSM, all fluorophores on the path of the illumination light are excited, while only the photons emitted from the light sheet waist region are collected and focused on the detection camera. Since the 3D imaging of the whole sample is realized by imaging subdivided sample volumes, a large number of subvolumes are illuminated repetitively before they are actually imaged, so these subvolumes could be bleached long before they are imaged. The degree of the photobleaching can be estimated by the ratio of the length of the excitation light sheet to length of the sample. Clearly, the smaller the ratio, the more severe the photobleaching, which limits either the highest spatial resolution that can be achieved to image a sample of a given size or the largest sample size that can be imaged at a required spatial resolution. A new 3D imaging strategy seems to be necessary to solve the problem.

Tiling light sheet microscopy is more efficient than conventional LSM to image cleared tissues with high spatial resolution because the need of sample translations to image the entire sample is reduced by translating the light sheet instead of the sample, which is much faster, more accurate and flexible. The advantage is more pronounced with a larger detection FOV, which is limited by the Nyquist sampling limit required to reach the desired spatial resolution and the pixel number of the detection camera. Therefore, the imaging efficiency of the microscope can be further improved by using a detection camera with more pixels or a detection camera array (Fan et al.,2019).

In summary, we developed a versatile tiling light sheet microscope with an advanced multicolor 3D imaging ability to image cleared tissues in 3D with micron-scale to nanometer-scale spatial resolution. The microscope is compatible with all tissue clearing methods and flexible to adapt to different applications. It is also reliable and easy to operate because of the ability to be aligned via phase modulation. Altogether, the microscope could make 3D cleared tissue imaging much more capable and feasible in biomedical research.

## Acknowledgements

We thank Ming Yin, Yi Sun, Kiryl D. Piatkevich, Yalin Wang, Xufeng Wu, and John Hammer for helpful discussions. We thank advanced biomedical technology core facility for the facility support and technical assistance. We thank the laboratory animal resource center for the facility support and technical assistance. L.G., J.M.J., S.C. and K.L. acknowledge the support from the Westlake University. L.G. acknowledges the support from the Zhejiang Province Natural Science Foundation LR20C070002.

## Contributions

L.G. conceived the project and designed the microscope; L.G., J.M.J., and S.C. designed the experiments and supervised the research; L.G., X.L.L. and Y.W. constructed the microscope; L.G. and Y.W. developed the software; Y.C., D.Z., C.W., X.Z.L., R.F., H.X, X.S.Z., X.Y., B.Z., Y.Y.C. and Y.F. prepared the samples; L.G., X.L.L. and Y.C. performed the imaging experiments with the tiling light sheet microscope; D.D.Z and X.Z.L. performed the imaging experiments with the Zeiss Z1 light sheet microscope. L.G., J.M.J., S.C., K.L., X.L.L., D.Z., C.W., and B.C.C. analyzed the data; L.G., J.M.J., S.C. and K.L. wrote the paper with the input of all authors.

## Materials and Methods

### Microscope configuration

CW laser beams with excitation wavelengths of 488 nm and 561 nm (Coherent, OBIS 488 nm, OBIS 561 nm) are combined and expanded to a 1/e^2^ beam diameter of ~7 mm (L1=30 mm, L2=250 mm) and sent to a binary spatial light modulator (SLM) assembly. The binary SLM assembly consists of a polarizing beam splitter cube, a half-wave plate and a 1280×1024 binary SLM (Froth Dimension Displays, SXGA-3DM) to modulate the illumination light. The modulated light is focused on an optical slit to block the undesired diffraction light and further conjugated to a galvanometer (Cambridge Technology, 6215H 7mm mirror) through relay lenses (L3=300 mm, L4=175 mm). The Galvo mirror is used to direct the illumination light to one of the two symmetrical illumination paths by offsetting the initial angle of the Galvo mirror and to create a virtual excitation light sheet to illuminate the sample by scanning the excitation laser beam. The laser beam is further conjugated to the real pupils of both excitation objectives (Mitutoyo MY5X-802 or MY5X-803) through two pairs of relay lenses (L5=L7=150 mm, L8=L9=250 mm) and illuminate the sample from two opposite directions. MY5X-803 has a 0.28 NA compared to the NA of MY5X-802, so that thinner light sheets can be generated by using MY5X-803.

The emission fluorescence is collected with the detection objective (Olympus MVPLAPO 1X, Olympus MVPLAPO 2XC, Nikon CFI90 20XC Glyc or Nikon N40X-NIR) installed on a modified Olympus MVX10 Macro Zoom microscope and imaged onto a sCMOS camera (Hamamatsu, Orca Flash 4.0 v3) through a filter wheel. A MVX10 microscope is used in the detection path for several reasons. First, the microscope is equipped with multiple flatness corrected, long WD objectives specifically designed to image large samples. Second, the microscope is capable to adjust the magnification from 0.63× to 6.3×, so that the size of the FOV can be adjusted quickly. Third, it has a conventional microscope design so that the operation of the microscope is more convenient and familiar to most scientists.

The imaging chamber subsystem consists of an imaging chamber, a 3D sample translational stage, a buffer tray and a sample holder (Figure 1f-g). The sample is mounted on the sample holder driven by the 3D translational stage that consists three single axis stages (PI Q-522.230 or PI Q-545.240). The sample is immersed in the imaging buffer during imaging. The buffer tray is used as both the mounting base of the imaging chamber and a container of the spilled imaging buffer. The imaging chamber is mounted with a magmatic kinematic mount so that imaging chambers containing different imaging buffers can be switched and placed in the same position. Multiple sample mounting methods are used to mount samples of different sizes, shapes and mechanical strengths (figure supplement 7). The sample holder is attached to the sample translational stage via two high pull magnets to enable fast sample loading.

The microscope control system (figure supplement 16) consists of a computer workstation, a NI PCIe-6323 DAQ card and two NI SCB-68A (National Instrument) connector blocks. The control system receives input parameters from the control software developed in LabView and then generates synchronized control signals to control the opto-mechanical and opto-electrical devices of the microscope. The control software is also used to calibrate the microscope and generate modulation phase maps to operate the microscope in different imaging modes. The major microscope components and their costs are listed in Table supplement 1.

### Microscope Calibration

The microscope needs to be calibrated before imaging. Briefly, the modulation phase can be expressed as (*A*_1_ × *Tilt x*_*p*_ + *A*_2_*Tilt x*_*b*_) + *Tilt y* + *Defocus*, which can also be expressed using Zernike Polynomials as (*A*_1_ × *Z*(1, −1)*x*_*p*_ + *A*_2_ × *Z*(1, −1)*x*_*b*_) + *Z*(1,1) + *Z*(2,0). *A*_1_ is the area on the SLM corresponding to the intensity profile of the illumination light at the excitation objective pupil plane. It is used to adjust the geometry of the excitation beam. *Tilt x*_*p*_ is applied to the incident illumination light on *A*_1_ area of the SLM. It allows the modulated illumination light to pass the optical slit. *Tilt x*_*b*_ is applied to the incident light on the rest area, *A*_2_, of the SLM, so that the abandoned light is blocked by the optical slit. *Tilt y* is applied to the illumination light to adjust the axial position of the excitation beam along the detection optical axis to keep the excitation light sheet in focus. *Defocus* is applied to the illumination light to tile the excitation beam along the excitation light propagation direction.

To calibrate the microscope, the imaging chamber is filled with the imaging buffer used to image the sample. The excitation beam can usually be visualized on the camera using the autofluorescence of the imaging buffer. *A*_1_ is determined at first based on the desired spatial resolution and number of tiles. *Defocus*_*left*_ and *Tilt y*_*left*_ are determined in the next by tiling the excitation beam on the left side of the FOV and moving it in focus. *Defocus*_*right*_ and *Tilt y*_*right*_ are determined afterwards in the same way. The microscope is calibrated after these parameters are determined. The phase maps used to operate the microscope are generated via interpolation using the calibrated parameters. For instance, the modulation phase to tile the excitation beam in the middle of the FOV is (*A*_1_ × *Tilt x*_*p*_ + *A*_2_*Tilt x*_*b*_) + (*Tilt y*_*left*_ + *Tilt y*_*right*_/2 + (*Defocus*_*left*_ + *Defocus*_*right*_/2. The generated phase map is binarized and shifted to correct the beam tilt before being applied to the SLM.

### Image analysis

The acquired image data are analyzed as follows (Figure supplement 17).

1. Raw images acquired from different subvolumes are analyzed individually following the same procedure as follows.
2. Raw images acquired in each subvolume are separated into multiple 3D image stacks that are corresponding to different tiling positions.
3. The region corresponding to the light sheet waist position in each separated 3D image stack is determined. An amplitude mask is generated for each separated image stack.
4. The final 3D image of the subvolume is obtained by summing the products of all separated 3D image stacks and the corresponding amplitude mask determined in the previous step.
5. The same procedure is repeated to obtain the 3D images of all subvolumes. The amplitude masks obtained in the third step are usually the same for all subvolume. They can also be adjusted for each individual subvolume if necessary.
6. The 3D image of the entire sample volume is obtained by registering and merging the reconstructed 3D images of all subvolumes using Amira. The 3D images of all subvolumes usually need to be down sampled before being registered and merged due to the large data size.

### Sample preparation

#### CUBIC

CUBIC was performed as previously described (Susaki et al., 2014; Susaki et al., 2015). The mice were deeply anesthetized using pentobarbital (~150 mg/kg of body weight) and perfused with heparinized 0.1 M PBS (10 U/ml of Heparin) for 5-10 minutes at room temperature until the blood was washed out, followed by 4% paraformaldehyde (PFA) in 0.1 M PBS (pH 7.4). Subsequently, organs from these animals were dissected out and post-fixed in 4% PFA overnight at 4 °C, and washed with 0.1 M PBS for 2 hours twice at room temperature and delipidated with CUBIC reagent 1 (25 wt% urea, 25 wt% Quadrol and 15 wt% Triton X-100). PI (5 μg/ml) was added to reagent 1 for nuclear stained specimens. Replace the reagent every 2 days until organs were sufficiently cleared. After delipidation, organs were washed with PBS for 2 hours three times at room temperature with gentle shaking, and then immersed in CUBIC-2 (25 wt% urea, 50 wt% sucrose and 10 wt% triethanolamine) at 37 °C with gentle shaking overnight. The reagent was replaced with fresh reagent afterwards and organs were further incubated until transparent.

#### PEGASOS

Dissected mouse tissues and bones were immersed in 5 ml of 4% paraformaldehyde (PFA) on a roller-mixer at room temperature for 24 hours and washed in PBS three times afterwards. Bones were decalcified next in 20% Ethylenediaminetetraacetic acid(EDTA) for 4 days, while soft tissues were directly immersed in a 25% N,N,N′,N′-Tetrakis (2-Hydroxypropylethylenediamine; Quadrol) water solution for 2 days and 5% ammonium solutions for 1 day in sequence for decolorization. Samples were immersed in tert-butanol (tB) solutions with gradient concentrations of 30% v/v, 50% v/v, and 70% v/v, for 1 to 2 days for delipidation, then dehydrated with 70% tB v/v, 27% polyethylene glycol (PEG) v/v, 3% Quadrol v/v for 2 days. Samples were finally cleared in BB-PEG medium composed of 75% benzyl benzoate (BB), 22% PEG, and 3% Quadrol for 2 days. Cleared tissues were preserved in BB-PEG at room temperature after clearing.

#### Adipo-Clear

The immunostaining and clearing of the mouse mammary gland, human breast cancer and paracancerous tissues (Approved by the independent ethics committee of Zhejiang cancer hospital (approval No: IRB-2015-176) were performed with a modified Adipo-Clear method.

Briefly, the tissues were fixed with 4% PFA overnight, permeabilized with 1% Triton-X100-PBS for 1day, and incubated with primary and then secondary antibodies in 37C° for 4 days respectively. Rabbit anti-keratin 14 (Biolegend, Poly19053, 1:500) and rat anti-CD200 (Biolegend, 123802, 1:200) or mouse anti-CD146 (ebioscience, 11-1469-41, 1:100) were used as primary antibodies. Alexa Fluor 488 or 594 conjugated donkey IgGs (Jackson) are used as secondary antibodies. After immunostaining, the tissues are dehydrated through 25%, 50%, 75% and 100% methanol, then delipidated with dichloromethane (DCM, sigma, 270997) and finally incubated in dibenzyl ether (DBE, sigma, 108014) before imaging.

#### MAP (mouse tissue)

Tissue expansion of mouse tissue with MAP was performed as previously described (Ku et al., 2016). After anesthesia, the mice were first perfused with a mixture of 5% acrylamide (AA), 0.05% bis-acrylamide (BA), and 0.8% sodium acrylate (SA) in PBS, followed by perfusion with a mixture of 4% PFA, 30% AA, 0.1% BA, 10% SA, 0.1% VA-044 in PBS. The brain was dissected out and incubated in 40 mL of the same fixative solution at 4 °C for 3 days and then at room temperature (RT) for another 3 days with gentle shaking. Subsequently, the brain was incubated for 2 hours at 45°C using EasyGel (LifeCanvas, EasyGel) for hydrogel tissue hybridization. Then, the brain hydrogel was cleared by incubating in a solution of 200 mM SDS, 200 mM NaCl, and 50 mM Tris in DI water (pH titrated to 9.0) and gently shaken at a gradient increasing temperature. Finally, the denatured brain was immersed in1000 mL DI water for 48 hours to allow the brain to expand. During DI water incubation, the solution was changed every 12 hours.

#### MAP (Planarian Schmidtea mediterranea)

Asexual planarians were starved for 1 week before fixation. Planarians were dipped in a 1x PBS solution with 5% N-Acetyl Cystein (NAC) for 5 mins at room temperature (RT), then fixed with 4%PFA for 2 hours, followed by washing with 10 ml of PBS/0.01% (wt/vol) sodium azide for at least 15 mins twice at room temperature (RT). Immerse the sample in 500 μl tissue clearing reagent mixed with 5μg/ml PI for 12 hours. The planarians were immersed into a mixture of 30% AA, 0.1% BA, 10% SA, 0.1% VA-044 in PBS at 4 °C for 12 hours with gentle shaking and then at 37 °C for 0.5 hour. Finally, the denatured worms were incubated in DI-water for 2 hours at RT for tissue expansion.

### Imaging conditions

The imaging conditions of the imaged samples are listed in Table supplement 2.

