## supplementary information for "A versatile tiling light sheet microscope for cleared tissues imaging"

Video 1. 3D rendering of a Thy1-YFP mouse brain. Associated with Figure 2.

Video 2. Part 1: 3D renderings of a SMACre::Ai14 mouse brain. Associated with Figure 3a. Part 2: 3D renderings of a subvolume of the imaged sample. Associated with Figure 3d. Part 3: 3D rendering of blood vessel traces within the imaged sample. Associated with Figure 2j.

Video 3. 3D rendering of a dual color mouse immune-stained mammary gland. Associated with Figure 4a.

Video 4. 3D renderings of two similar sample volumes in two Thy1-eGFP mouse brains cleared with PEGASOS and CUBIC. Associated with Figure 5.

Video 5. 3D renderings of a $9\times11\times5$ $mm^3$ sample volume in an expanded Thy1-eGFP mouse brain. Associated with Figure 6.

Video 6. Comparison of the same sample volume in an expanded Thy1-eGFP mouse brain imaged with a submicron spatial resolution and a nanometer-scale spatial resolution. Associated with Figure 7a.

Video 7. Part 1: 3D renderings of a sample volume in an expanded CAG-zsGreen mouse brain imaged using a discontinuous light sheet with a submicron spatial resolution. Part 2: Comparison of the same sample volume imaged with a submicron spatial resolution and a nanometer-scale spatial resolution. Associated with Figure 7k.

Video 8. 3D renderings of an expanded planarian Schmidtea mediterranea labeling stem cells and all cell nuclei imaged with a submicron spatial resolution and a nanometer-scale spatial resolution. Associated with Figure 8.

Download link:

https://westlakeu-my.sharepoint.com/:f:/g/personal/gaoliang_westlake_edu_cn/EvfqH0ESI2JLpGGOvHxFzngBoKKw4qh917XsvFQw8vmr_w?e=0l4ZTa

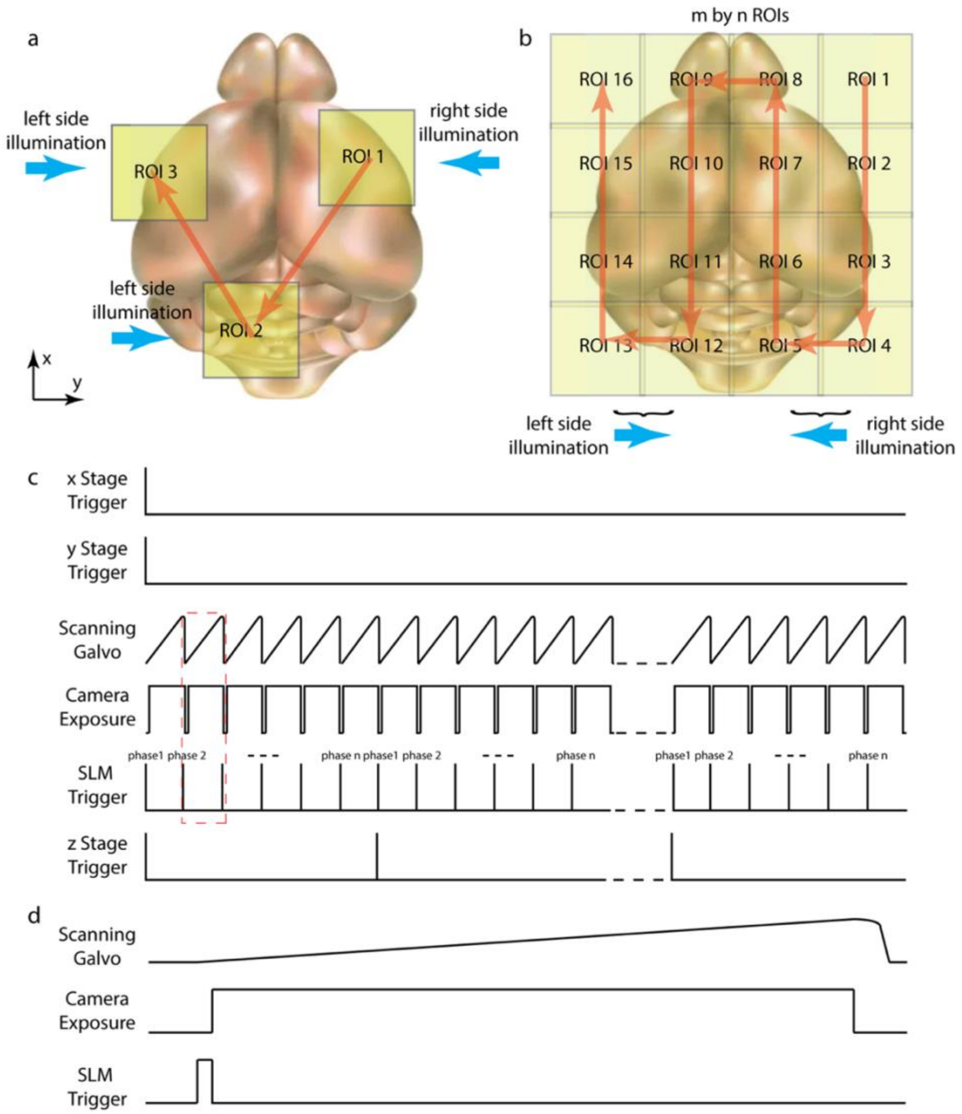


Figure supplement 1. 3D imaging modes of the microscope and the corresponding control signal sequences. (a) The sample is translated to image the preselected ROIs in 3D sequentially. The illumination light is sent from the assigned excitation objective. (b) A large sample volume is imaged by imaging an array of subvolumes in sequence. (c) Synchronized control signal sequences used to image an individual ROI in 3D with tiling light sheets. The sample is translated in lateral directions to position the target ROI in the detection FOV. The galvanometer, detection camera, SLM, and axial translational stage are synchronized to image the ROI in 3D by tiling the excitation light sheet at *n* positions in each image plane. (d) The synchronization of the scanning galvanometer, detection camera, and spatial light modulator in one camera exposure period marked in (c).

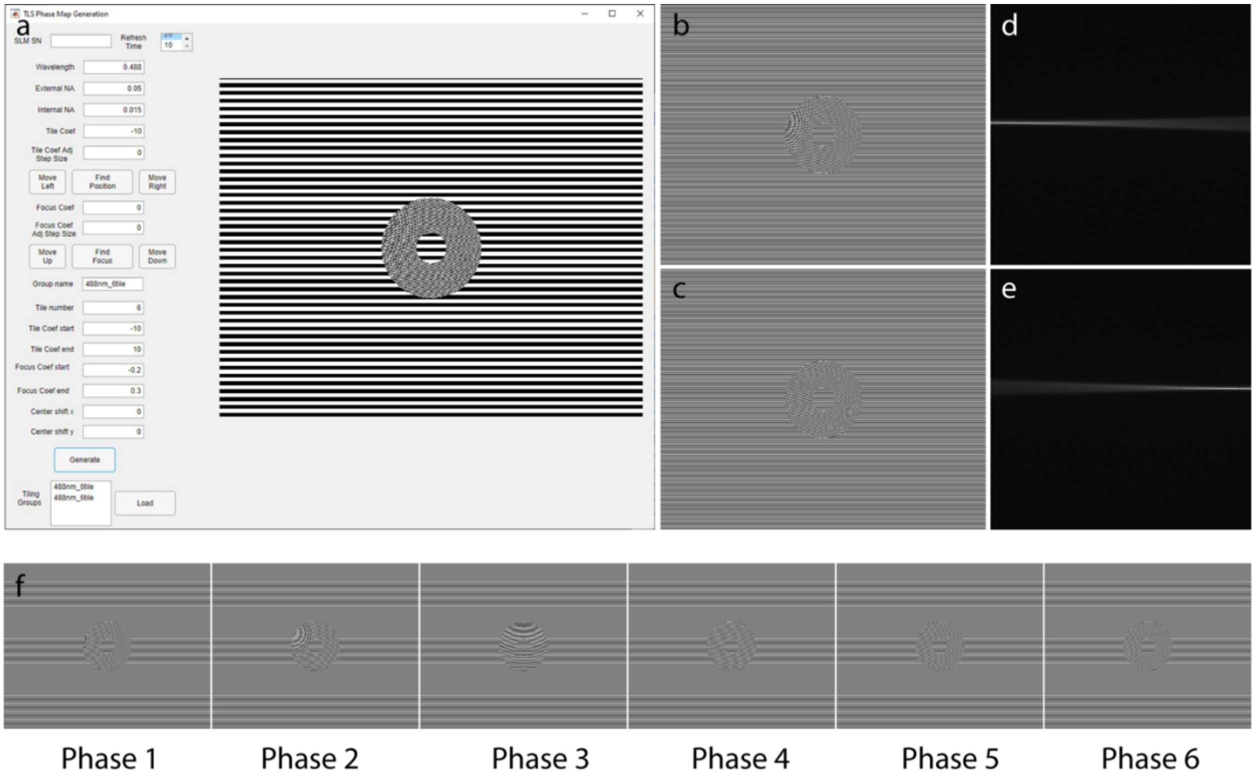


Figure supplement 2. Generation of the modulation phase maps. (a) The interface of the control software used to calibrate the microscope and generate the modulation phase maps. (b,c) Calibrated phase maps to tile the excitation light sheet on the left and right side of the FOV and keep the light sheet in focus at both positions. (c,d) The excitation beams generated by applying the phase maps in (b) and (c). (f) A group of phase maps interpolated with the calibrated parameters to tile the excitation light sheet at six positions in the FOV and keep the light sheet in focus at all positions.

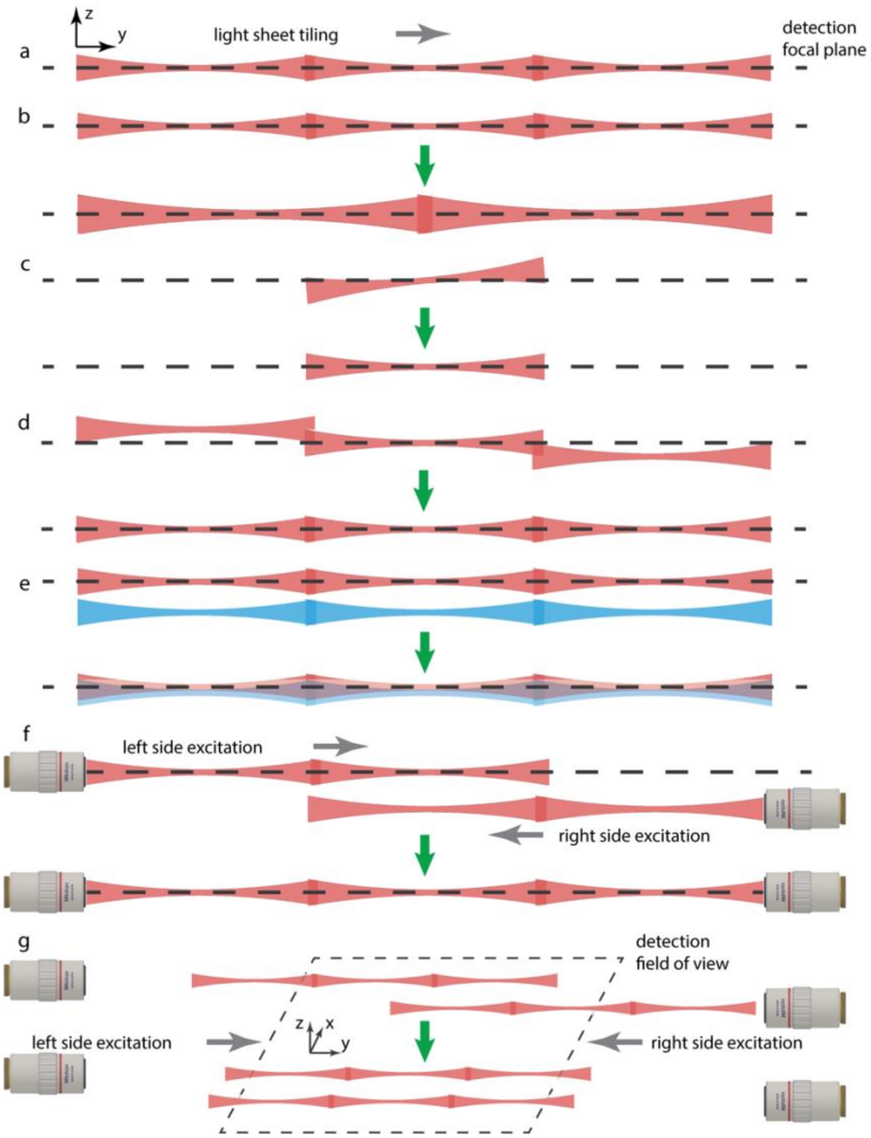


Figure supplement 3. The functions realized and the alignment errors corrected by modulating the phase of the illumination light via the binary SLM. (a) Tile a light sheet along the illumination light propagation direction in the image plane. (b) Change the geometry of the excitation light sheet. (c) Correct the tilt of the excitation light sheet. (d) Keep the excitation light sheet in focus at different tiling positions. (e) Keep the colinear alignment of the excitation light sheets of different excitation wavelengths. (f) Keep the excitation light sheets sent from both excitation objectives in focus. (g) Correct the lateral drifting of tiling light sheets sent from both excitation objectives.

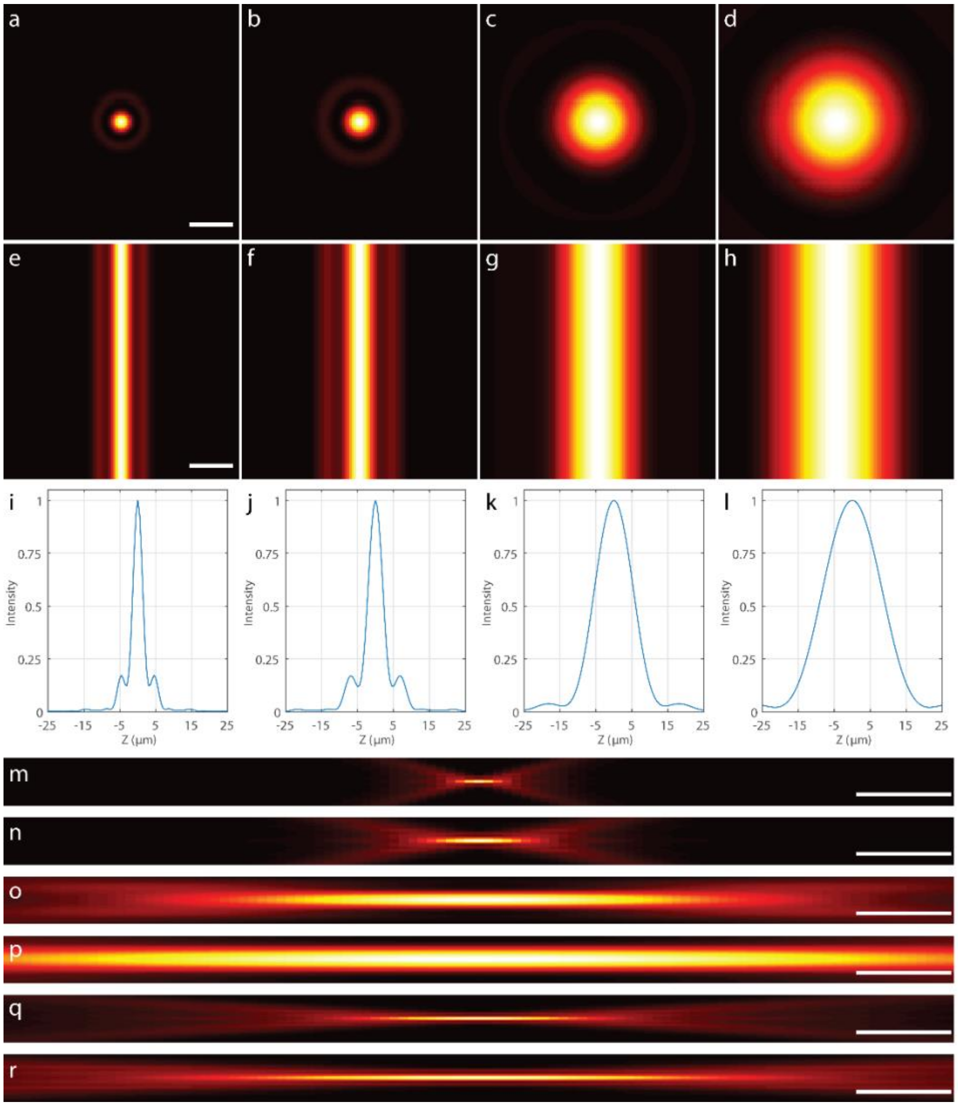


Figure supplement 4. Comparison of different excitation light sheets. (a-d) The lateral intensity profiles of the excitation beams created with excitation (numerical apertures) NAs of (a) $NA_{OD}$=0.08, $NA_{ID}$=0.03. (b) $NA_{OD}$=0.05, $NA_{ID}$=0.02. (c) $NA_{OD}$=0.02, $NA_{ID}$=0. (d) $NA_{OD}$=0.013, $NA_{ID}$=0. (e-h) The intensity profiles of the corresponding excitation light sheets obtained by scanning the excitation beams in (a-d). (i-l) The intensity plots of the excitation light sheets in (e-h). (m-p) The axial intensity profiles of the excitation light sheets in (e-h) showing the trade-off between the length and thickness of the excitation light sheet. (q,r) Zoom in views of (m) and (n). Scale bars, 10 μm (a,e), 1 mm (m-p), 200 μm (q,r).

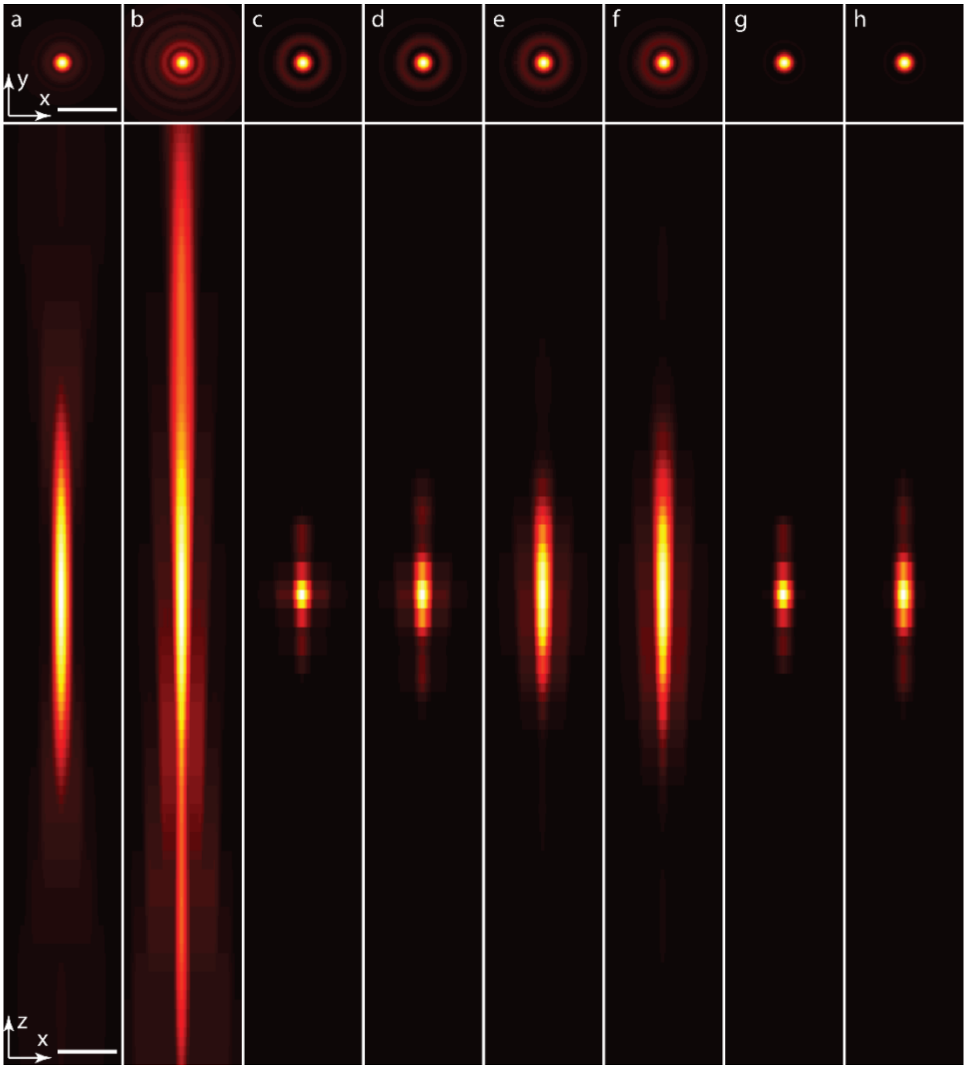


Figure supplement 5. Simulated point spread function (PSF) of the microscope obtained with a 0.25 numerical aperture (NA), 60 mm working distance (WD), air objective and the excitation light sheets in figure supplement 6e-h. (a) The ideal detection PSF. (b) The aberrated detection PSF with the point source located 10 mm below the surface of the imaging buffer with 1.5 refractive index (RI). (c-f) The final PSF obtained with the aberrated detection PSF and the excitation light sheets in figure supplement 6e-h. (g-h) The final PSF obtained with the ideal detection PSF and the excitation light sheets in figure supplement 6e-f. The final PSF obtained with the 0.25 NA air objective and a thin excitation light sheet is nearly the same as the ideal PSF due to the thin light sheet thickness. Scale bar, 5 µm.

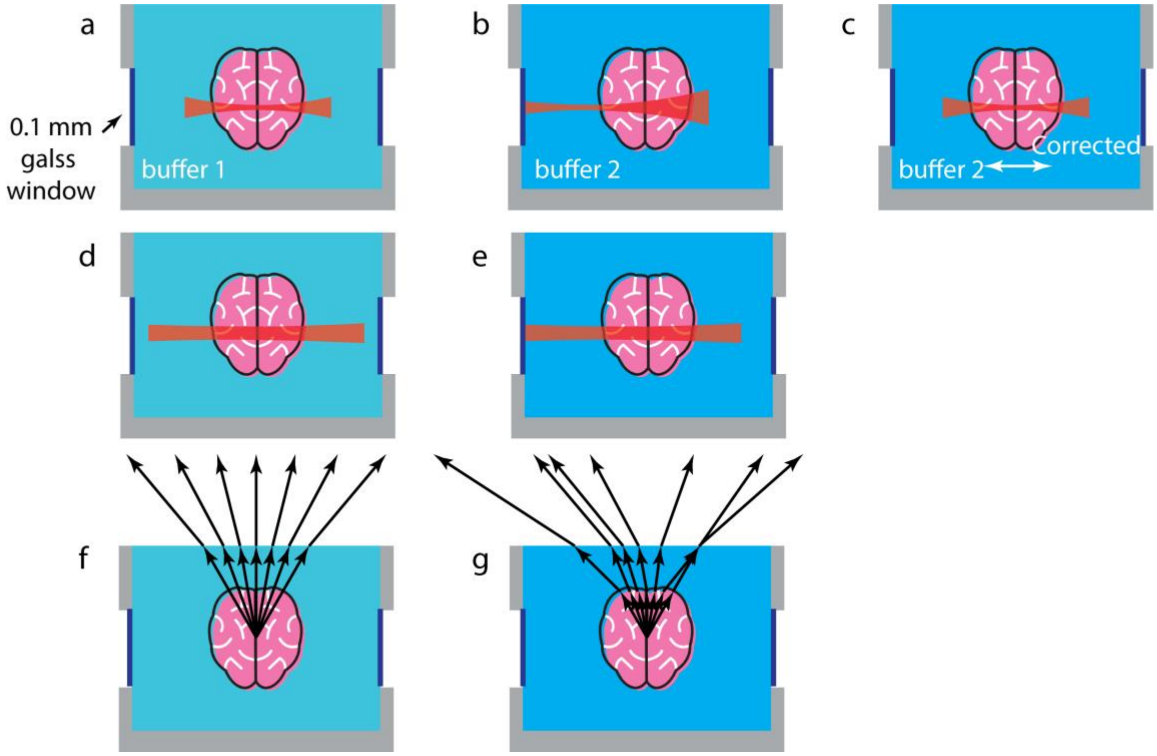


Figure supplement 6. The influences of the imaging buffer RI variation on the excitation light sheet and the RI mismatch on the emission fluorescence. (a-b) The RI variation of the imaging buffer causes lateral drift of the excitation light sheet. (c) The lateral drift of the excitation light sheet must be corrected in order to center the along the light propagation direction light sheet in the FOV. Otherwise, the spatial resolution decreases quickly due to fast diverging of the excitation light sheet. (d-e) A longer but thicker light sheet that is less affected by lateral drifting must be used to maintain a uniform spatial resolution in the FOV for conventional light sheet microscopes without the ability to correct the lateral drift, which decreases the axial resolution. (f) High-NA detection objectives are more vulnerable to the RI mismatch between air and the imaging buffer. (g) The RI mismatch between the imaging buffer and the sample could introduce strong optical aberrations due to the irregular sample surface.

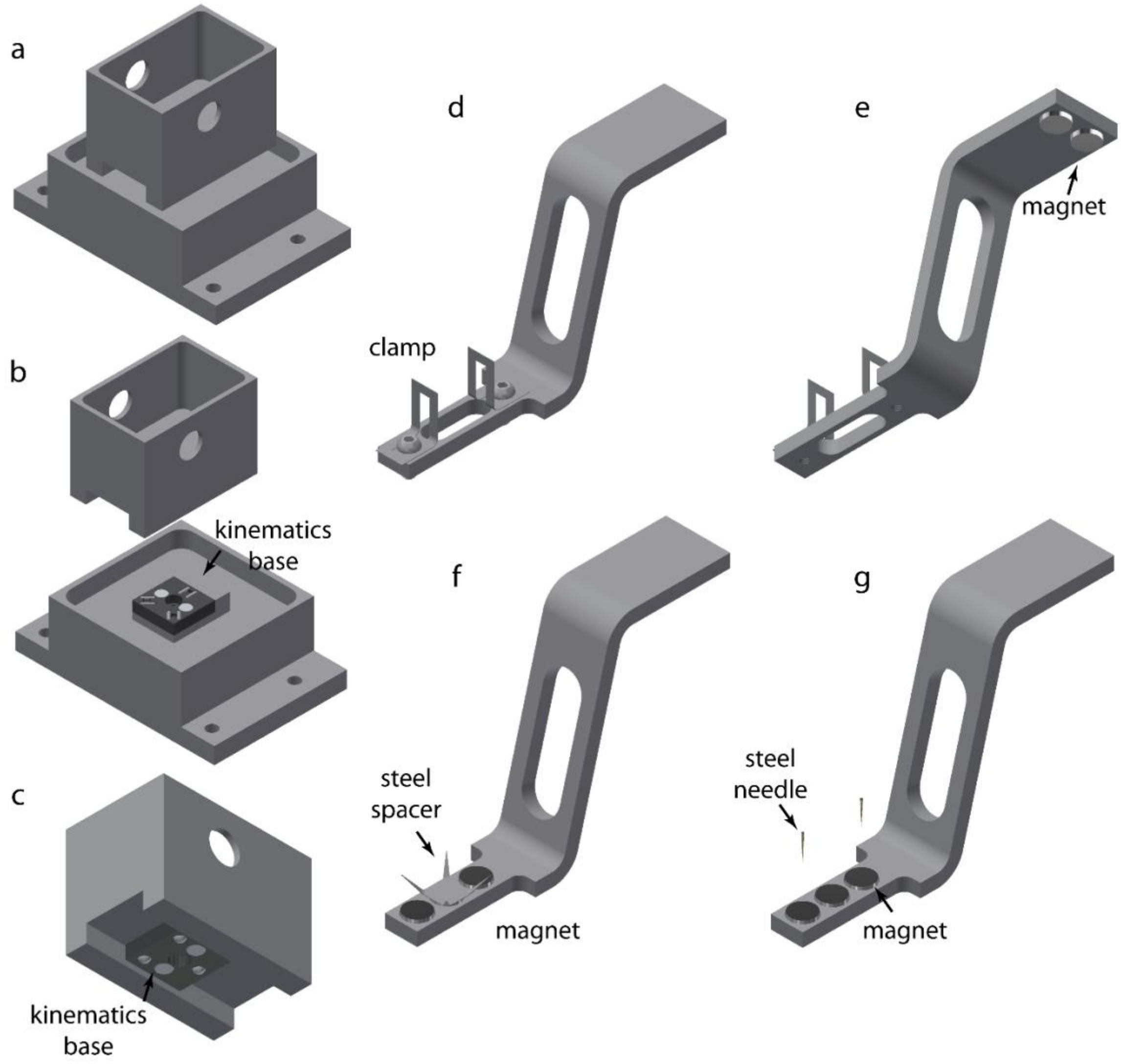


Figure supplement 7. The imaging chamber and sample holders of the microscope. (a-c) Imaging chambers containing different imaging buffer can be switched quickly by using a magnetic kinematic mount. (d-g) Different sample holders are used to mount samples of different sizes, shapes, and mechanical strengths. The sample holder is attached to the 3D translational stage via a pair of high pull magnets. (d,e) Both soft and stiff samples with more isotropic dimensions can be mounted with a clamp, or (f) glued on a steel spacer attached to the magnetic base. Different spacers can be used to fit samples of different shapes. (g) Soft and thin samples can be pinned on the magnets of the sample holder with a pair of steel needles.

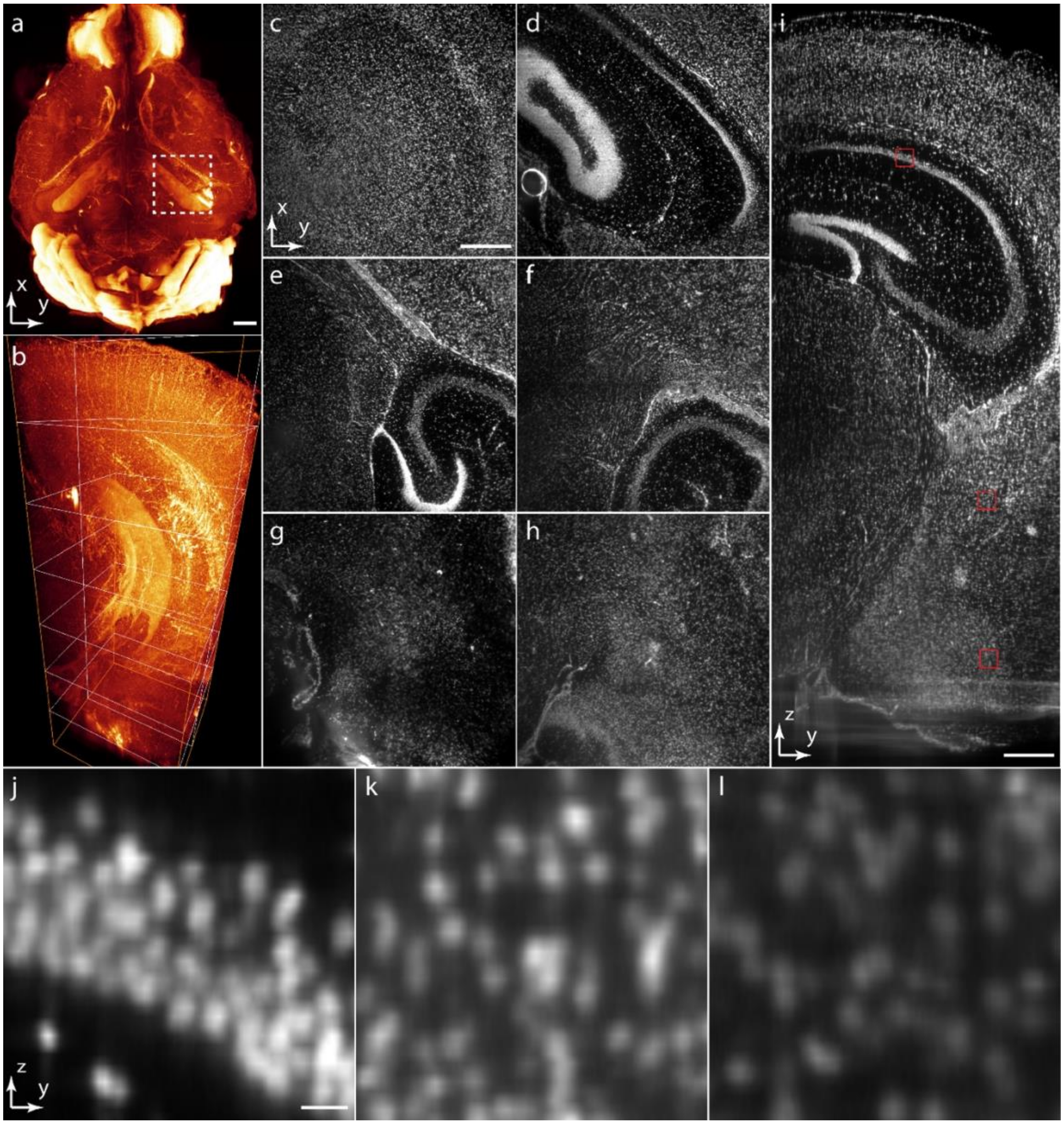


Figure supplement 8. (a) 3D rendering of a PI stained mouse brain cleared by CUBIC. (b) 3D rendering of the selected 2.6×2.6×7 $mm^3$ volume in (a). (c-h) Lateral image slices through the indicated planes in (b) at the depth from 1 to 6 mm below the sample surface. (I) Axial image slice through the indicated plane in (b). (j-l) Zoom in views of the selected 150×150 $\mu m^2$ areas at different depths in (i). Scale bars, 1 mm (a), 500 µm (c,i), 20 µm (j).

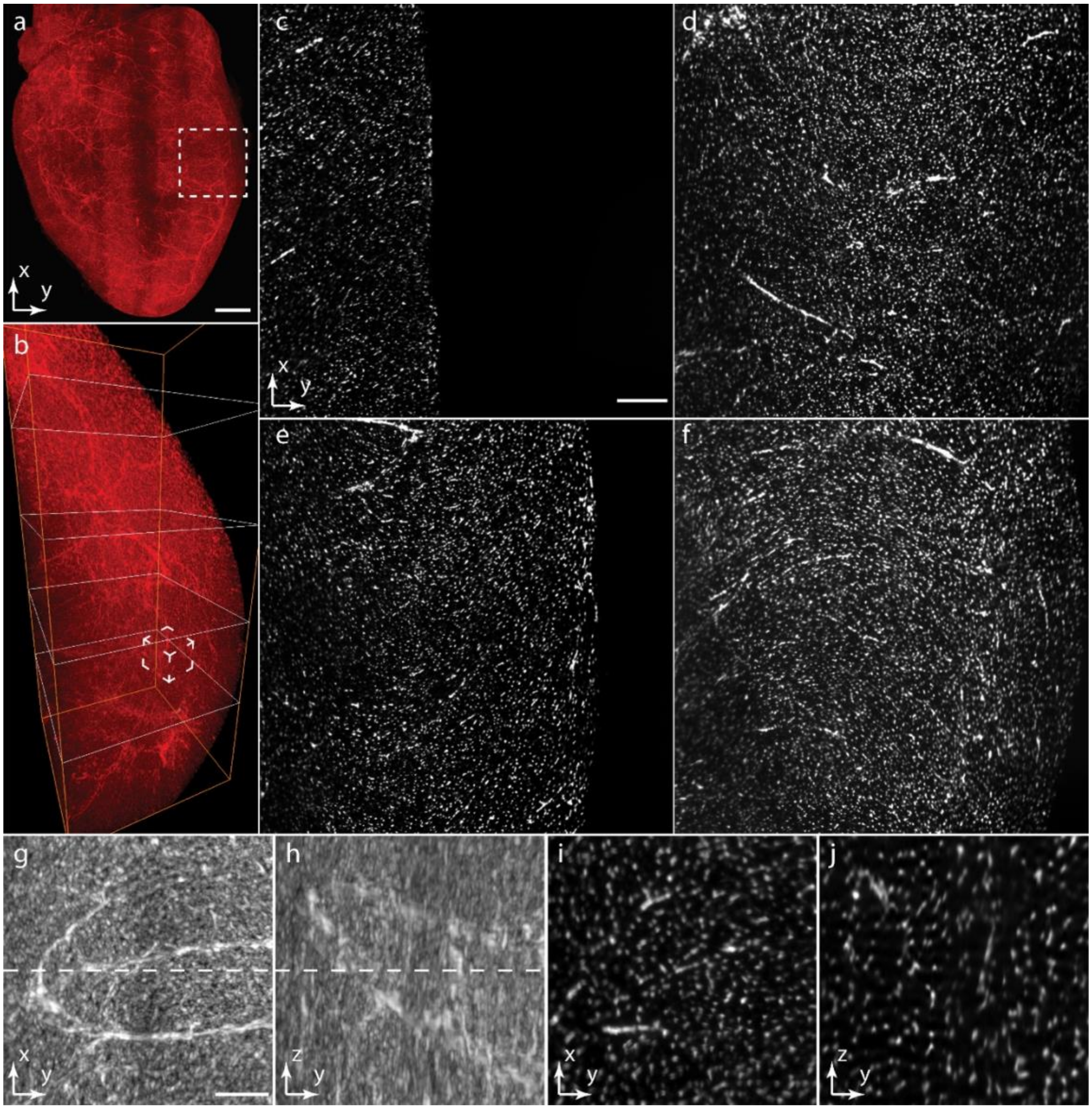


Figure supplement 9. (a) 3D rendering of a PI stained mouse heart cleared with CUBIC. (b) 3D rendering of the selected 2×2×5 mm$^3$ region in (a). (c-f) Lateral image slices through the indicated planes in (b) at the 1 to 4 mm depth. (g-x) Lateral and axial maximum intensity projections (MIP) of the selected region in (b). (I,j) Lateral and axial image slices through the indicated planes in (h) and (g). Scale bars, 1 mm (a), 200 µm (c), 100 µm (g).

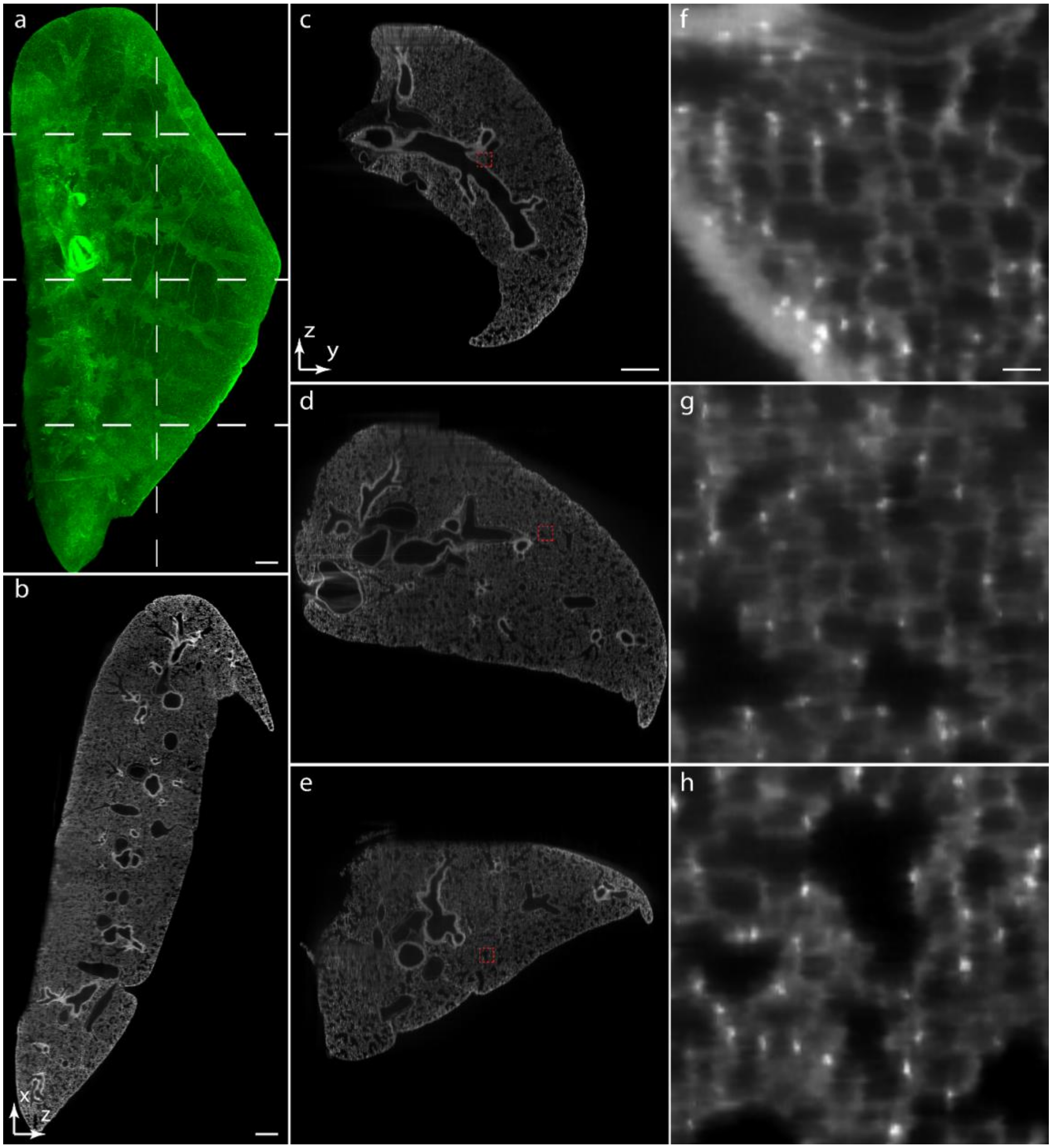


Figure supplement 10. (a) 3D rendering of a mouse lung cleared by PEGASOS. (b) Axial image slice of the mouse lung through the indicated planes in (a). (c-e) Axial image slices of the mouse lung through the indicated planes in (a). (f-h) Zoom in views of the selected 200 µm by 200 µm areas in (c-e). Scale bars, 500 µm (a-c), 20 µm (f).

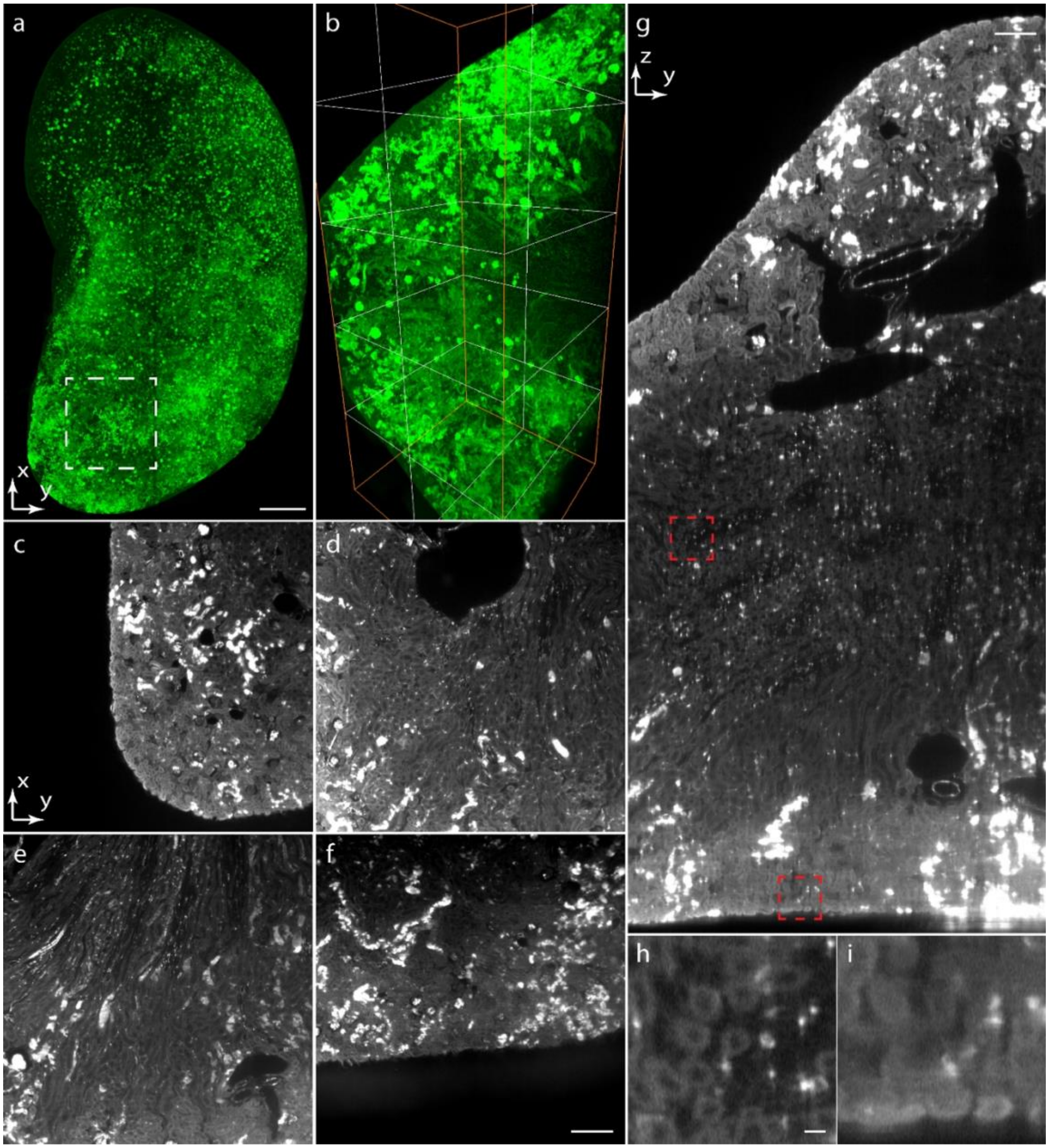


Figure supplement 11. (a) 3D rendering of a mouse kidney cleared with PEGASOS. (b) 3D rendering of the selected 2×2×5 $mm^3$ volume in (a). (c-f) Lateral image slices through the indicated planes in (b) at the depth 1 to 4 mm depth. (g) Lateral image slice through the indicated plane in (b). (h,i) Zoom in views of the selected 200 µm by 200 µm areas at different depths in (g). Scale bars, 1 mm (a), 200 µm (f,g), 20 µm (h).

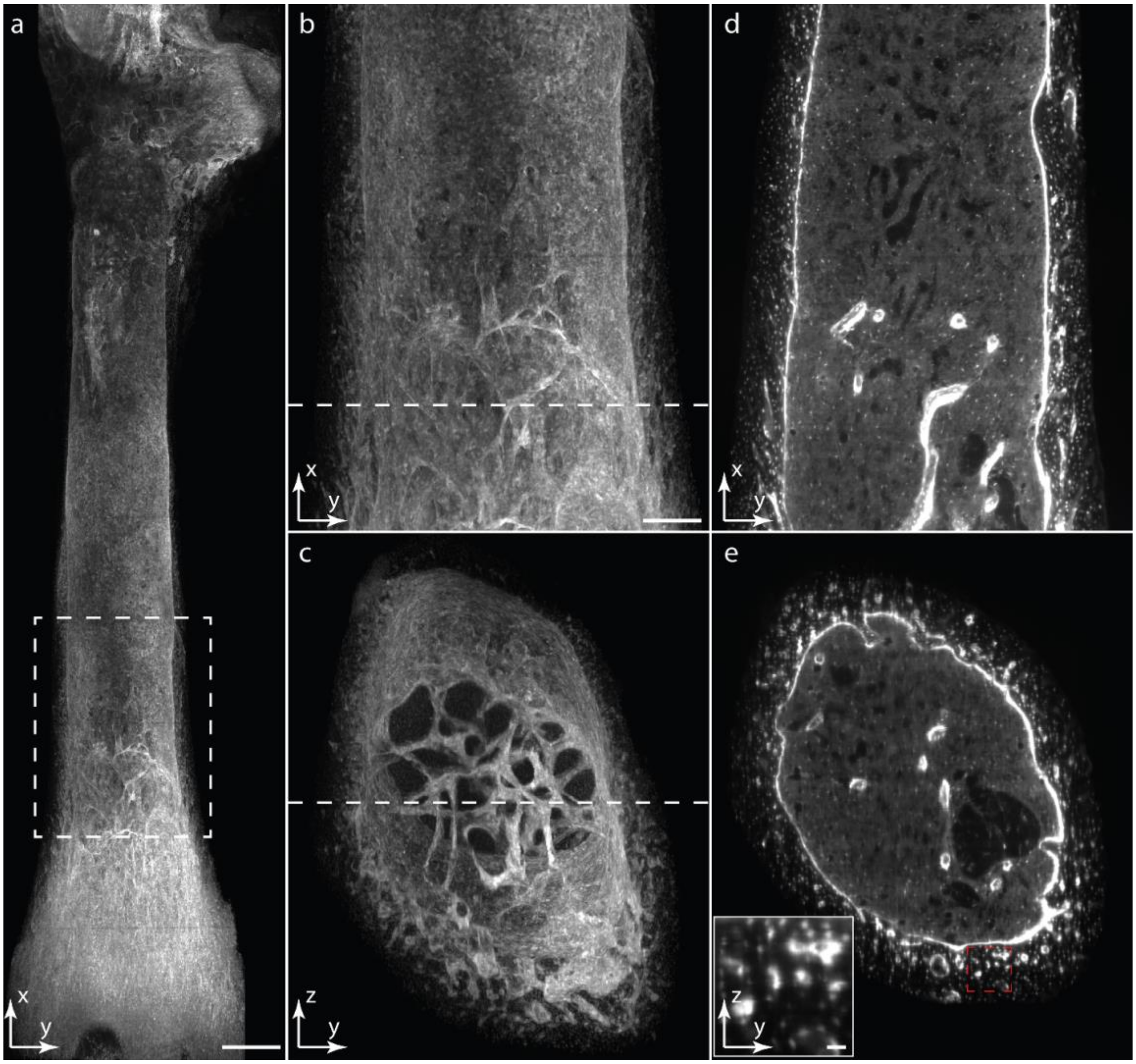


Figure supplement 12. (a) 3D rendering of a mouse femur cleared by PEGASOS. (b,c) Lateral and AXIAL MIPs of the selected region in (a). (d,e) Lateral and axial image slices through the indicated planes in (c) and (b). The insert shows the zoom in view of the selected area in (e). Scale bars, 500 µm (a), 200 µm (b), 20 µm (e) insert.

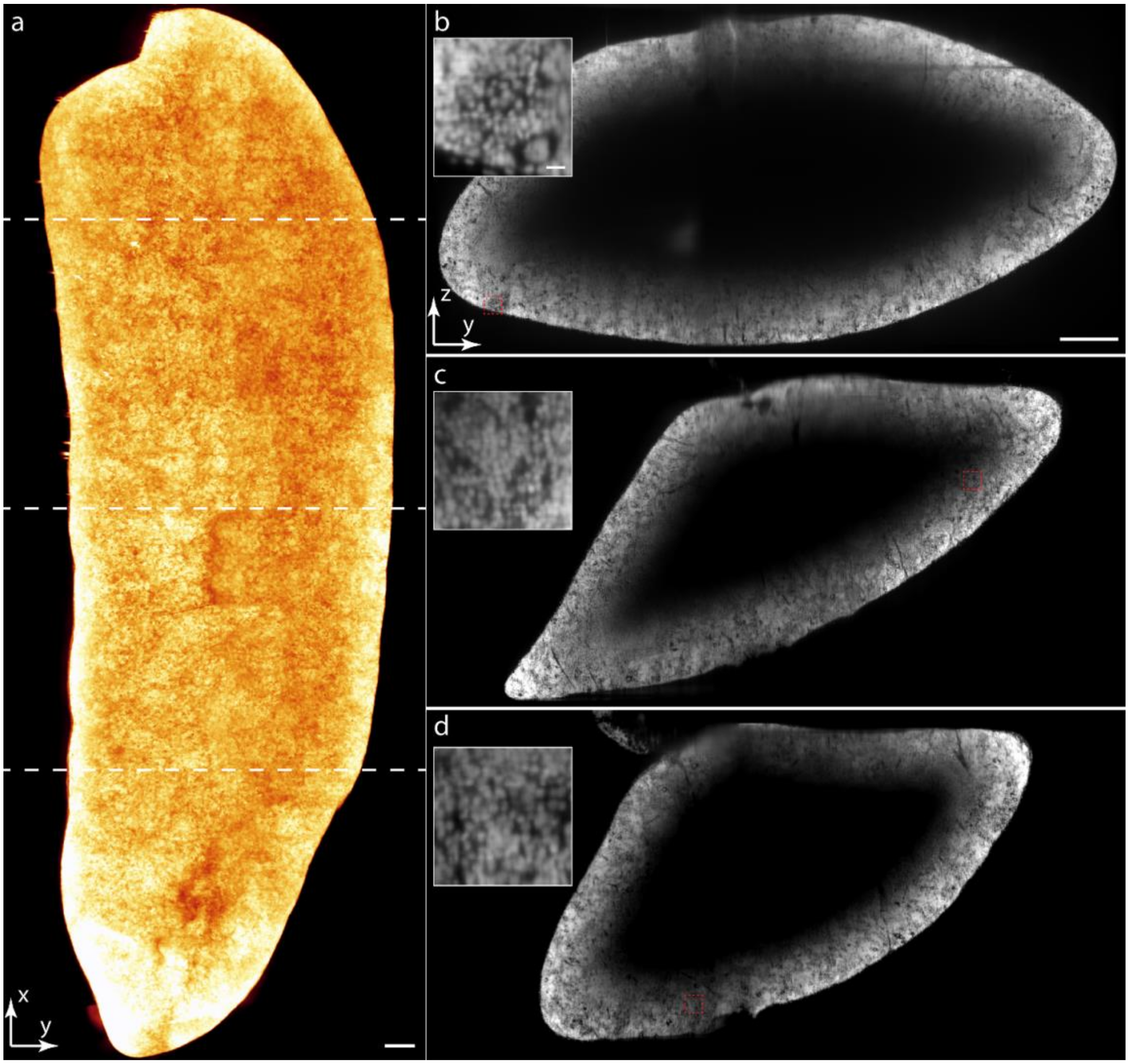


Figure supplement 13. (a) 3D rendering of a PI stained mouse spleen cleared by CUBIC. (b-c) Axial image slices of the mouse spleen through the marked planes in (a). The inserts show zoom in views of the selected 150×150 μm$^2$ areas in (c-d). The result shows the extremely dense cellular structure of mouse spleen, that perhaps prevented the PI stain from penetrating to the center of the organ. Scale bars, 500 μm in (a,b), 20 μm (b) insert.

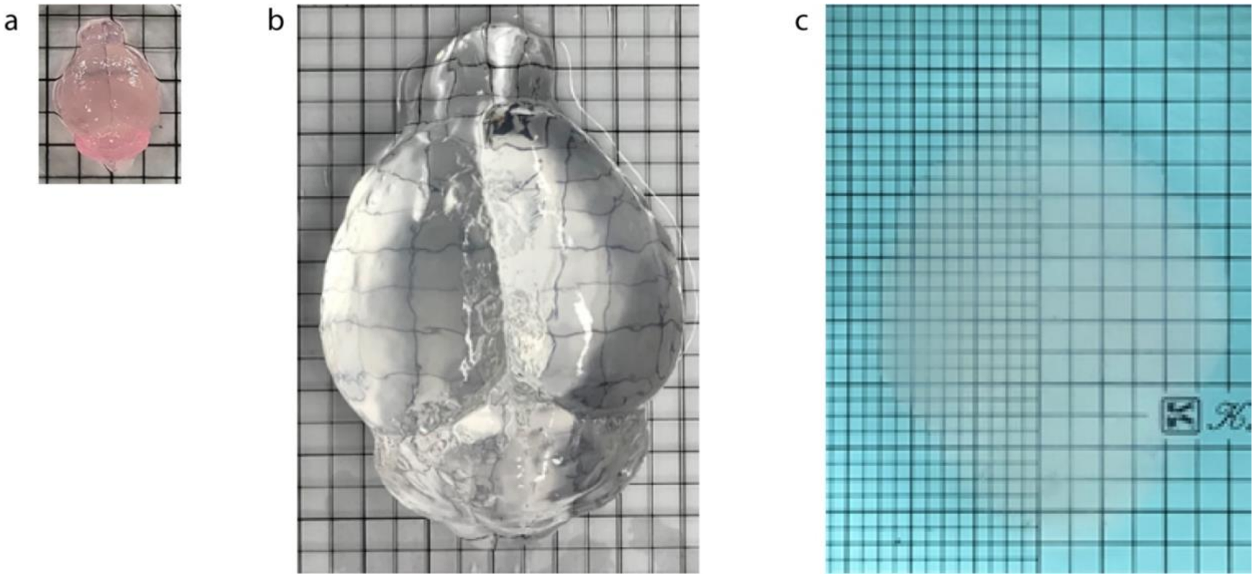


Figure supplement 14. A mouse brain expanded with MAP. (a) Before expansion. (b) After expansion. (c) The expanded brain immersed in water. Blue color dye was added into the water to enhance the photo contrast.

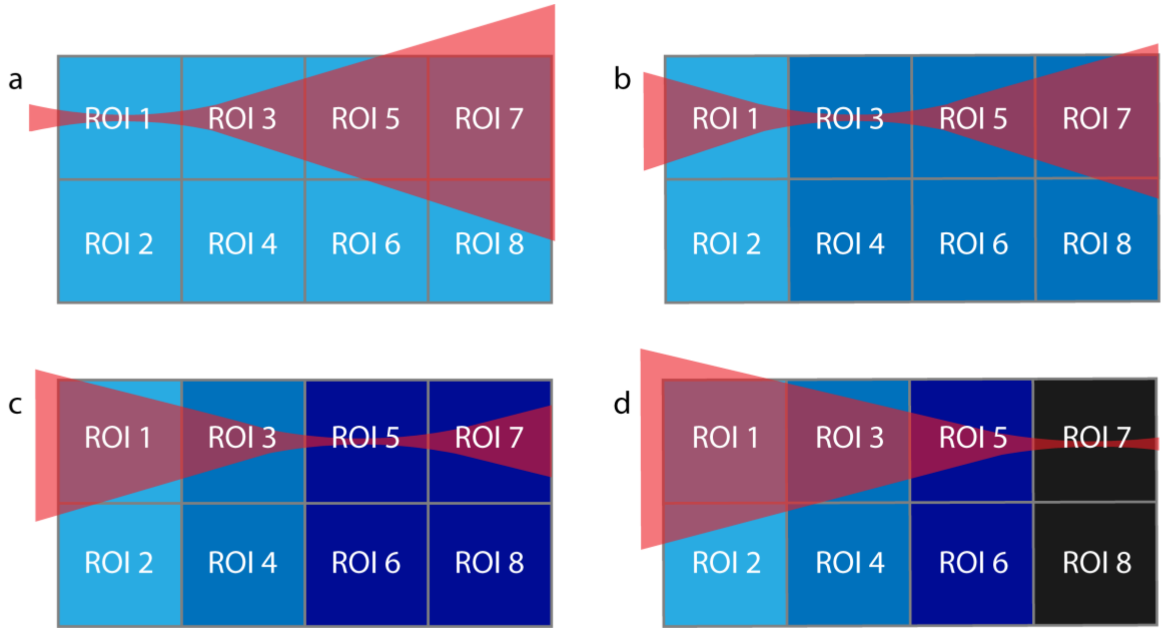


Figure supplement 15. Photobleaching of a large cleared tissue when it is imaged with LSM. A large sample is imaged by imaging subvolumes from ROI 1 to ROI 8 in sequence. (a) ROI 3-ROI 8 are also illuminated when ROI 1 and ROI 2 are imaged. (b) ROI 5-ROI 8 are also illuminated when ROI 3 and ROI 4 are imaged. (c) ROI 7 and ROI 8 are also illuminated when ROI 5 and ROI 6 are imaged. (d) ROI 7 and ROI 8 are photobleached before they are imaged.

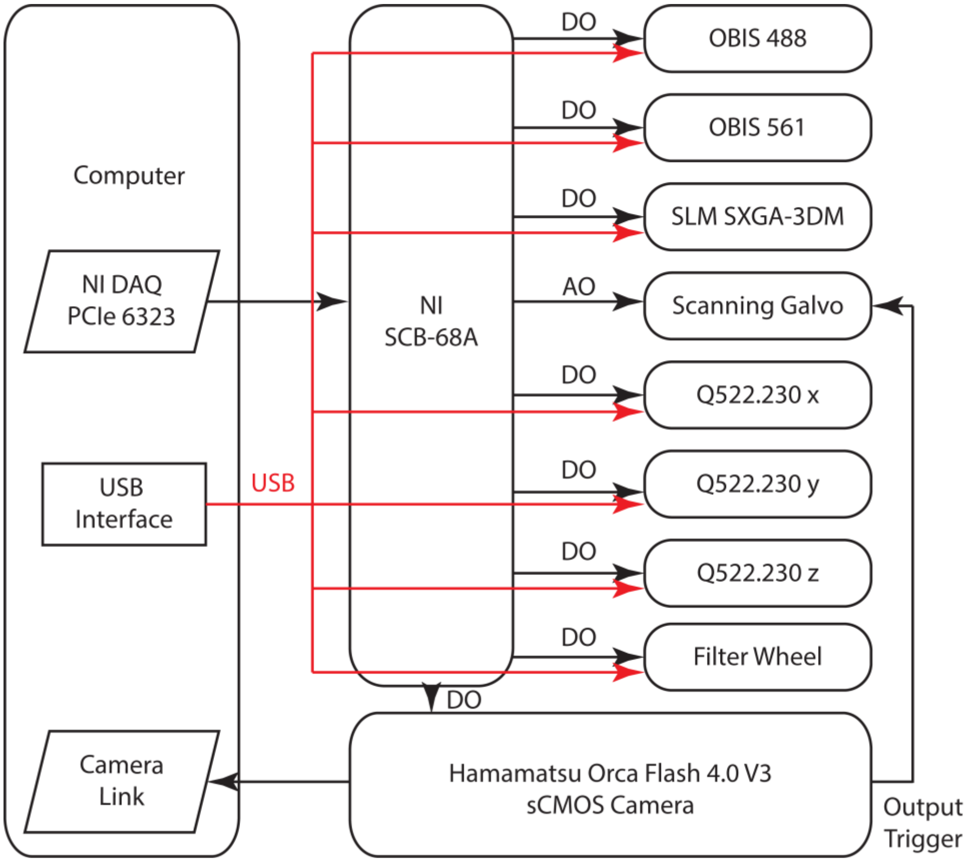


Figure supplement 16. The control system diagram. The control system consists of a computer workstation, a NI PCIe-6323 DAQ card and two SCB-68A connector blocks. Synchronized control signals are generated to control the opto-mechanical and opto-electrical devices of the microscope, which include a 150 mW OBIS 488 laser, a 150 mW OBIS 561 laser, a binary spatial light modulator (SLM), a galvanometer, a 3D translational stage consists of three identical Q522.230 linear stages, a filter wheel and a sCMOS camera. More excitation lasers can be added into the microscope for multicolor imaging if needed.

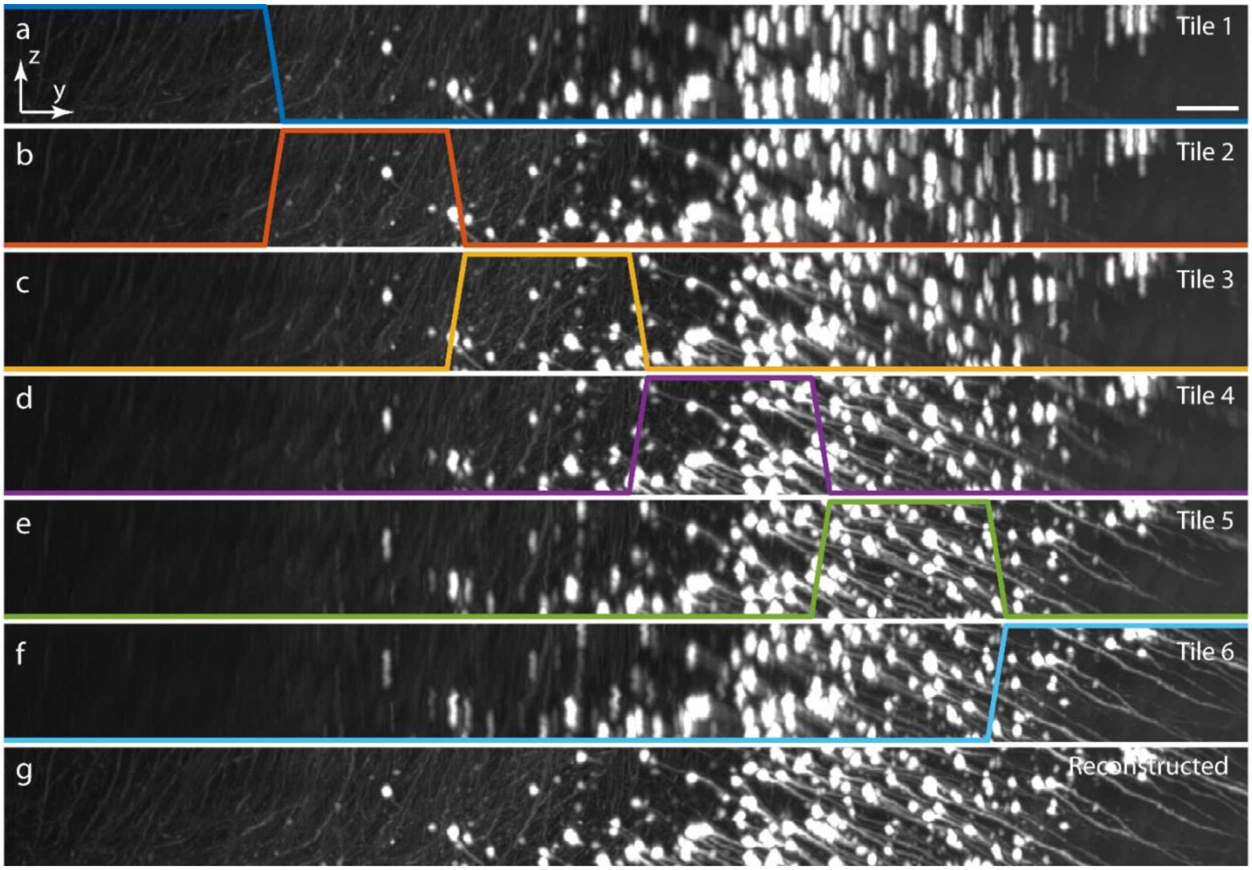


Figure supplement 17. Image reconstruction. (a-f) Axial MIPs of the separated image stacks collected from the same sample volume by tiling the excitation light sheets at six tiling positions. (g) Axial MIP of the reconstructed 3D image stack obtained by summing (a-f) multiplied by the corresponding amplitude masks.

Table supplement 1. The major components of the microscope.

| Components | Model | Cost (USD) |
|---|---|---|
| 488 nm laser | Coherent OBIS 488 LS 150 mW | 1,0000 |
| 561 nm laser | Coherent OBIS 561 LS 150 mW | 11,000 |
| Spatial light modulator | Forth Dimension Displays SXGA-3DM | 4,500 |
| Galvanometer | Cambridge Technology 6215H with 7 mm mirror | 1,500 |
| Excitation objective | Mitutoyo MY5X-802<br>0.14 NA, 34 mm WD | 1,000 × 2 |
| Optional excitation objective | Mitutoyo MY5X-803<br>0.28 NA, 34 mm WD | 1,000 × 2 |
| 3D translational stage | Physik Instrumente PI Q522.230 and controller | 3,000 × 3 |
| Detection subsystem | Olympus MVX10 with the associated components and detection objectives<br>MVPLAPO 1X, MVPLAPO 2X | 10,000 |
| Optional detection objective | Nikon CFI90 20XC Glyc | 20,000 |
| Optional detection objective | Nikon N40X-NIR | 3,000 |
| Detection camera | Hamamatsu Orca Flash 4.0 v3 | 15,000 |
| Other Optical and Optomechanical components | Thorlabs, Edmundoptics, Semrock, Bolder Vision | 30,000 |
| Machining | Customized | 3,000 |
| Control system | National Instruments PCIe-6323, 2×SCB-68A | 2,000 |
| Computer workstation | Customized | 7,000 |
| | | Basic configuration: 105,000<br>Complete configuration: 130,000 |

Table supplement 2. The sample information and imaging conditions.

| Sample | Clearing method | Labeling method | Sample size X×Y×Z (mm), ROI number × ROI size X×Y×Z (mm) | Excitation NA, (OD,ID) Tiles per FOV | Detection NA | Spatial resolution X×Y×Z (μm) | Imaging time/speed per channel |
|---|---|---|---|---|---|---|---|
| Figure 2 six-tile mode | PEGASOS | Thy1-YFP | 9.2×9.3×6 30×(2×2×6) | 0.05, 0.02 6 | 0.25 | 2×2×5 | 5 hours 30 s/mm³ |
| Figure 2 non-tile mode | | | 9.2×9.3×6 12×(4×4×6) | 0.015, 0 1 | 0.25 | 4×4×10 | 12 mins 0.6 s/mm³ |
| Figure 2 Zeiss Z1 | | | 9.2×9.3×6 72×(1.6×1.6×6) | 10 µm thick light sheet | 0.16 | 2.3×2.3×15 | 1.5 hours 10 s/mm³ |
| Figure 3a | PEGASOS | Transgenic SMACre::Ai14 | 8.7×8.5×6 28×(2×2×6) | 0.05, 0.02 6 | 0.25 | 2×2×5 | 5 hours 30 s/mm³ |
| Figure 3r | CUBIC | PI staining | 16.9×13.1×9 31×(2.6×2.6×9) | 0.04, 0.01 4 | 0.25 | 2.6×2.6×6 | 5 hours 10 s/mm³ |
| Figure 3s | CUBIC | PI staining | 9.6×8.5×7 26×(2×2×7) | 0.05, 0.02 6 | 0.25 | 2×2×5 | 4 hours 30 s/mm³ |
| Figure 3t | PEGASOS | Transgenic Pdgfrb-cre/tdTomato | 12.8×6.2×6 14×(2×2×6) | 0.05, 0.02 6 | 0.25 | 2×2×5 | 3 hours 30 s/mm³ |
| Figure 3u | PEGASOS | Transgenic Pdgfrb-cre/tdTomato | 11.7×7.6×5 25×(2×2×5) | 0.05, 0.02 6 | 0.25 | 2×2×5 | 4 hours 30 s/mm³ |
| Figure 3v | PEGASOS | Transgenic Pdgfrb-cre/tdTomato | 9×2.5×4 5×(2×2×4) | 0.05, 0.02 6 | 0.25 | 2×2×5 | 50 mins 30 s/mm³ |
| Figure 3w | CUBIC | PI staining | 18.5×7.9×4 38×(2×2×4) | 0.05, 0.02 6 | 0.25 | 2×2×5 | 5 hours 30 s/mm³ |
| Figure 4a | Adipo-Clear | Krt14 antibody staining Cd200 antibody staining | 12.6×8.2×3 27×(2×2×3) | 0.05, 0.02 6 | 0.25 | 2×2×5 | 3 hours/color 30 s/mm³ |
| Figure 4z | Adipo-Clear | Krt14 antibody staining Cd146 antibody staining | 12×7×4, 30×(2×2×4) | 0.05, 0.02 6 | 0.25 | 2×2×5 | 4 hours/color 30 s/mm³ |
| Figure 4dd | Adipo-Clear | Krt14 antibody staining Cd146 antibody staining | 8×6×5 12×(2×2×5) | 0.05, 0.02 6 | 0.25 | 2×2×5 | 2 hours/color 30 s/mm³ |
| Figure 5a | PEGASOS | Thy1-eGFP | 0.3×1.1×1.5 4×(0.3×0.3×1.5) | 0.12, 0.04 4 | 1.0 | 0.3×0.3×1.5 | 26 mins 0.8 hours/mm³ |
| Figure 5h | CUBIC | Thy1-eGFP | 0.3×0.3×1.5 | 0.12, 0.04 4 | 1.0 | 0.3×0.3×1.5 | 6.5 mins 0.8 hours/mm³ |

| | | | | | | | |
|---|---|---|---|---|---|---|---|
| Figure 6 | MAP | Thy1-eGFP | 9×11×5<br>30×(2×2×5) | 0.05, 0.02<br>6 | 0.25 | 2×2×5<br>(0.4×0.4×1<br>with<br>expansion) | 4 hours<br>25 s/mm³ |
| Figure 7a | MAP | Thy1-eGFP | 2.3×2.3×2 | 0.05, 0.02<br>6 | 0.25 | 2×2×5<br>(0.4×0.4×1<br>with<br>expansion) | 3.5 mins<br>20 s/mm³ |
| Figure 7d | MAP | Thy1-eGFP | 0.25×0.25×0.5 | 0.12, 0.04<br>4 | 0.8 | 0.33×0.33×1<br>(0.07×0.07×0.2<br>with<br>expansion) | 3.5 mins<br>1.8 hours/mm³ |
| Video 7 part1 | MAP | CAG-zsGreen | 3.8×3.8×7<br>4×(2×2×7) | 0.08, 0.03<br>3<br>Three-waist discontinuo us light sheet | 0.25 | 2×2×4<br>(0.4×0.4×0.8<br>with<br>expansion) | 28 mins<br>15 s/mm³ |
| Video 7 part2 | MAP | CAG-zsGreen | 1×1×1.1<br>25×(0.25×0.25×1.1) | 0.12, 0.04<br>4 | 0.8 | 0.33×0.33×1.5<br>(0.07×0.07×0.3<br>with<br>expansion) | 2 hours<br>1.2 hours/mm³ |
| Figure 7l | MAP | CAG-zsGreen | 0.25×0.25×0.6 | 0.12, 0.04<br>4 | 0.8 | 0.33×0.33×1.5<br>(0.07×0.07×0.3<br>with<br>expansion) | 3 mins<br>1.2 hours/mm³ |
| Figure 8<br>Submicron resolution | MAP | smedwi-1 antibody staining<br>PI staining | 4.3×9.3×4.7<br>10×(2.3×2.3×4.7) | 0.05, 0.02<br>6 | 0.25 | 2×2×5<br>(0.4×0.4×1<br>with<br>expansion) | 1.5 hours/color<br>20 s/mm³ |
| Figure<br>Super-resolution | MAP | smedwi-1 antibody staining<br>PI staining | 1×1×1<br>25×(0.25×0.25×1) | 0.12, 0.04<br>4 | 0.8 | 0.33×0.33×1<br>(0.07×0.07×0.2<br>with<br>expansion) | 2 hours/color<br>1.5 hours /mm³ |
